# A shared extracellular plasticity program drives brain repair and tumor progression

**DOI:** 10.64898/2026.08.13.744612

**Authors:** Juan Andrés Sánchez, Mariana B Santos, Ana Cristina Ojalvo-Sanz, Daniela Pão Alvo, Anabel R Simões, Sara Monteiro-Ferreira, Patricia Das Neves Borges, Tânia Carvalho, Adriana Sánchez-Danés, Sergio Casas-Tintó, Christa Rhiner

**Author notes:** Equal contributions (Juan Andrés Sánchez, Mariana B Santos).

## Abstract

Injury induces complex multicellular repair responses that restore tissue integrity and replenish lost cells. In the brain, regeneration is limited and depends on the proliferation of neural and glial progenitors. But how these cells remodel the microenvironment to support repair remains poorly understood. Here, we profile dividing cells in the injured fruit fly brain revealing signatures of neural stem cell activation and ECM remodeling. We find that rapid activation of a Heparan Sulfate-binding factor (Hpsl) and changes in glypicans condition the extracellular space for proliferation. Hpsl promotes signalling of scarce Dpp/BMP ligands in brain cells with latent proliferative capacity and regulates organismal resilience to brain injury. We further show that malignant cells in fly and mouse brains promote similar HS-interactions to support tumor progression. Together, our findings identify extracellular proteoglycan remodeling as a key regulator of brain repair and uncover a common plasticity program engaged by brain damage and tumorigenesis.

## Introduction

Tissue damage triggers extensive molecular and cellular changes that activate plasticity programs required for repair. Such responses include remodeling of the extracellular microenvironment, release of growth-promoting signals and proliferation of local quiescent cells, which collectively contribute to tissue repair and functional recovery (Gurtner et al., 2008; Eming et al., 2017). Repair processes typically strongly rely on re-activation of developmental programs (Fazilaty and Basler, 2023; Goldman and Poss, 2022). However, the extent to which these regenerative programs can be activated varies considerably across tissues (Tanaka and Reddien, 2011; Poss, 2010).

Among adult tissues, the brain presents a particularly challenging environment for proliferation with recent transcriptional studies detailing limited growth factor signalling and a shift towards inflammatory cues (Bormann et al., 2024; Han et al., 2024). While neurogenesis is sustained within specialized niches, or to a very low extent at the lesion site (Obernier and Alvarez-Bullya, 2019; Frisén, 2016; Magnusson et al., 2014), injury can also induce stem cell-like properties in astrocytes and trigger extensive state changes in glial populations (Sirko et al., 2013; Zamboni et al., 2020; Burda and Sofroniew, 2014; Burda et al., 2022), which influence CNS function due to intimately linked neuro-glial interactions. Although these responses are increasingly recognized as beneficial for acute repair (O’Shea et al., 2024; Frisén, 2016), the molecular programs that drive injury-induced proliferation and regenerative microenvironment remodeling remain poorly understood, particularly within the understudied extracellular space of the brain (Tønnesen et al., 2023).

Genetically tractable *Drosophila*, with a brain composed of functionally analogous neuronal and glial subtypes (Bitern et al., 2021), have been successfully used to elucidate crucial aspects of nervous system repair such as glial phagocytic activity and coordinated break-down of damaged neurons (Ziegenfuss et al., 2008; Osterloh et.al., 2012; Szabo et al., 2023), together with insulin-driven neuro-glial cross talk in the regenerative larval ventral nerve cored (VNC) (Harrison et al., 2021). Using a *Drosophila* model of brain injury, we previously found that stab lesions induce rapid proliferative response and the activation of normally quiescent Deadpan/HES1-like-expressing neural progenitor cells, that can form new neurons at the injury site (Simões et al., 2022; Li et al., 2020; Fernández-Hernández et al., 2013). Similar damaged-induced plasticity has also been observed in response to lesions in the central brain and the adult VNC (Kato et al., 2009; Crocker et al., 2021; Foo et al., 2017, Casas-Tintó et al., 2025).

The underlying molecular mechanisms mediating regenerative responses in the adult fly brain, revealed conserved pathways such as resilience-promoting AP1-activation in glia and neuronal JNK signalling (Byrns et al., 2021; Ayaz et al., 2008), Wnt/Swim-dependent activation of neural progenitors in response to hypoxia-sensing and pro-mitotic ROS release by glia (Simões et al., 2022; Alves et al., 2026). Nevertheless, the mechanisms by which proliferative cells drive brain plasticity and support tissue repair remain poorly understood.

Here we analyzed injury-induced changes specifically in dividing cells post injury and functionally delineate a multi-cellular signalling circuit that facilitates regeneration and confers resilience to brain injury. We find that rapid injury-driven remodeling of the extracellular space, particularly the proteoglycan landscape, promotes plasticity in the normally non-permissive adult fly brain by facilitating Dpp/BMP signalling in cells with latent proliferative capacity. Finally, we show that core components of this conserved core plasticity axis are also co-opted by invasive brain cancers in flies and mice, revealing molecular parallels between the regenerative microenvironment and tumor-malignant growth in the nervous system.

## Results

### Transcriptional hallmarks of regenerating cells after brain injury

Proliferative cells have been reported in the optic lobe (OL) of *Drosophila* following a stab lesion (Alves et al., 2026, Simões et al., 2022; Fernández-Hernández et al., 2013), making this an attractive model to study adult brain plasticity **(Figure 1A)**. Consistent with previous reports, we detected increased signals of the mitotic marker phospho-histone H3 (pH3) in adult OLs 72 hs after injury (AI), which was markedly reduced when the cell cycle regulator *cdk1* was specifically knocked-down by RNAi in Deadpan-expressing progenitor cells (*dpnT2A-GAL4*) or glial cells (*repo-GAL4*) in the adult brain with the GAL4/UAS and GAL80^ts^ expression system (**Figure 1B and S1A**). As expected, only a low pH3 signal was observed in non-injured OLs (**Figure S1A**).

**Figure 1.**
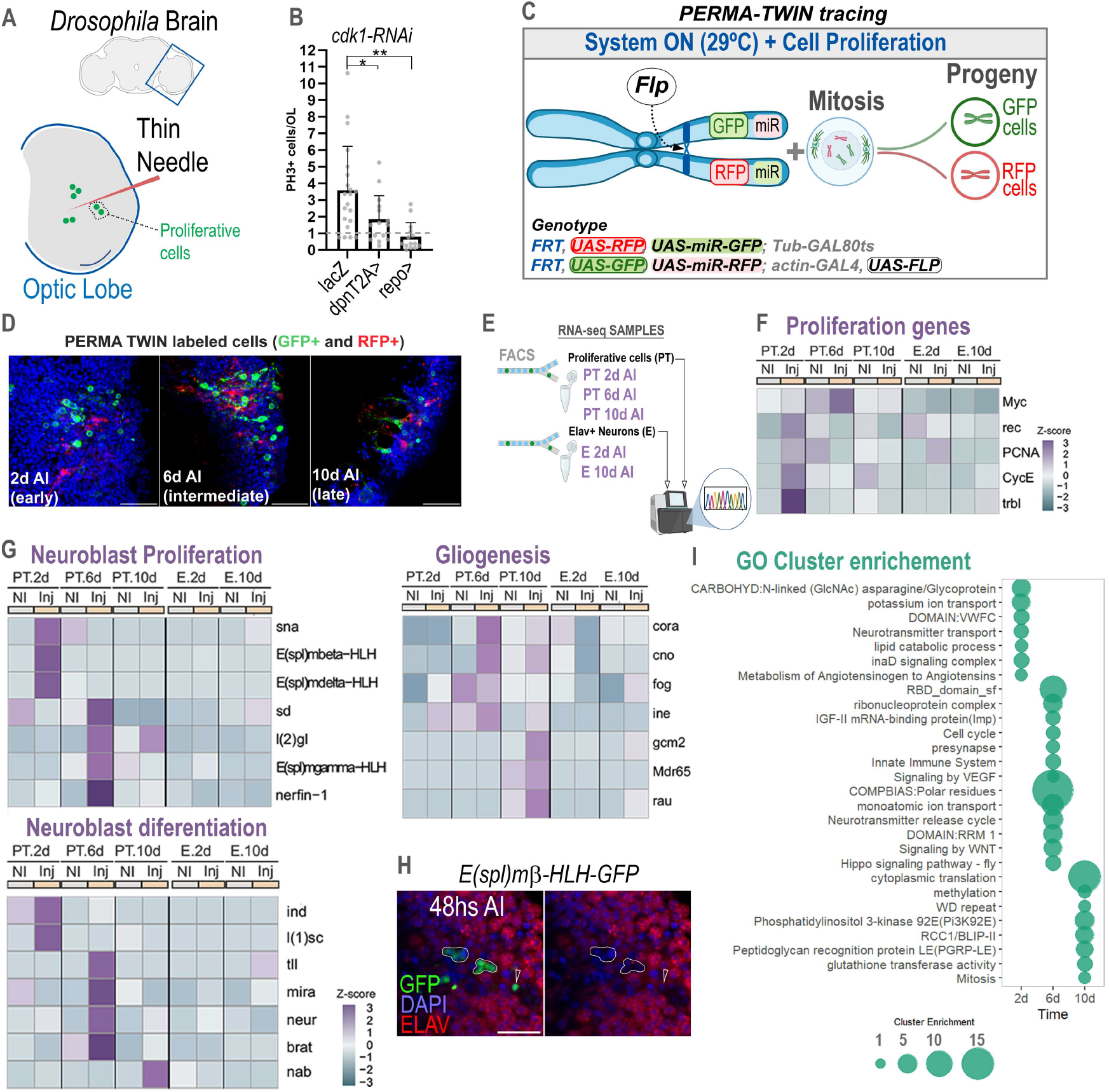
Transcriptional signature of injury-induced proliferative cells in the adult *Drosophila* optic lobe. (A) Schematic representation of the adult *Drosophila* brain highlighting the optic lobe (OL) injury model. An acute stab lesion was generated by introducing a needle into the OL, triggering a proliferative response. (B) Quantification of pH3+ mitotic cells. *cdk1* knockdown in glia (*repo>*) and progenitor (*dpnT2A>*) cells significantly reduced injury-induced proliferation. Data is shown as mean ± SD. *p <0.05, **p <0.01 as assessed by Kruskal-Wallis test. (C) Schematic of the PERMA-TWIN lineage-tracing strategy used to permanently label progeny derived from dividing cells. Mitotic recombination generates GFP+ and RFP+ daughter cell clones that allow identification and isolation of injury-responsive proliferative populations. (D) Representative images of PERMA-TWIN-labeled cells (GFP+,RFP+) in the optic lobe at 2, 6, and 10 days after injury (AI), corresponding to early, intermediate, and late stages of the regenerative response. Nuclei are counterstained with DAPI (blue). (E) Experimental design for transcriptomic profiling. PERMA-TWIN-labeled proliferative cells (PT) and Elav+ neuronal populations (E) were isolated by FACS from OLs 2,6 and 10 days after injury (AI) and from corresponding non-injured (NI) OLs (not shown) and subjected to RNA sequencing. (F) Heatmap showing expression dynamics of representative proliferation-associated genes in PT and Elav+ populations during the injury response. NI (no injury), Inj (injury). (G) Heatmaps showing temporal changes in genes associated with neuroblast proliferation, gliogenesis, and neuroblast differentiation following injury. (H) Representative image of E(spl)mβ-HLH-GFP expression 48 h AI. GFP marks reporter activity, ELAV labels neurons, and DAPI labels nuclei. (I) Gene Ontology enrichment analysis of differentially expressed genes in proliferative cells across the regenerative time course. Bubble size indicates enrichment score. Scale bars: 10 µm (D), (H)

To gain access to the proliferative cells, we activated a mitotic-dependent cell labelling system dubbed “perma-twin” in adult flies, which permanently labels cells after division with membrane GFP and RFP (**Figure 1C**) (Fernández-Hernández et al., 2013). We performed OL lesions to trigger proliferation and sorted GFP+ and RFP+ cells from non-injured and injured OLs by FACS and processed GFP+ cells that showed higher signal for RNAseq at 2, 6 and 10 days AI (**Figure 1D**). In addition, we collected GFP-expressing neurons (*elav*>GFP) to compare signatures in proliferating cells to injury-induced change in post-mitotic neurons (**Figure 1E**).

RNA-seq analysis followed by PCA revealed distinct clustering of perma-twin (PT) and Elav+ (E) cells, indicating cell-type–specific responses to injury (**Figure S1B**). Notably, PT cells displayed a high number of differentially expressed genes at 2 days (477 upregulated and 109 downregulated) when comparing injured versus non-injured conditions (**Figure S1C and Table 1**). At that time point, the mitotically marked cells showed activation of proliferation genes, including *CycE*, *PCNA*, and *rec* (**Figure 1F**), indicating a transient proliferative response. As expected, *myc* is also upregulated at both 2 and 6 days after injury, consistent with previous reports showing its essential role in driving cell divisions post injury (Fernández-Hernández et al., 2013; Simões et al., 2022).

To further characterize the regenerative cell population, we focused on genes associated with central nervous system development. Interestingly, we detected early enrichment (2d AI) of genes related to neuroblast proliferation, including members of the E(spl) complex (*E(spl)beta* and *E(spl)delta*), and at 6d AI, also *l(2)gl*, *E(spl)gamma* and *nerfin-1*, a gene required to maintain neuronal fate (Southall et al. 2014) (**Figure 1G**). At 6 and 10 days AI, we found increased expression of genes associated with neuronal differentiation (*brat*, *neur*, *nab*) and gliogenesis (*gcm2*, *cno*). Using GFP reporters, we confirmed the early induction (2dAI) of the HES transcription factors *E(spl)beta* in non-neuronal cells (Elav-) in line with the transcriptional profiling (**Figure 1H**). *E(spl)beta* activated in undifferentiated cells during early neurogenesis (Zacharioudaki et al, 2012), suggesting similar programs are turned on in dividing cells after injury.

We next compared the transcriptional signatures of mitotic cells across different time points following injury (**Figure S1B and Table 1**). Gene Ontology analysis revealed an early enrichment of extracellular matrix glycoproteins, potassium signalling components, and proteins containing von Willebrand factor type C domains commonly associated with ECM and BMP modulation (Zhang et al., 2007) (**Figure 1I and Table 2**). By 6 days post-injury, regenerating cells exhibited increased components of Wnt, Hippo, and VEGF signalling pathways, together with genes associated with presynaptic organization and neurotransmitter release. At later stages, transcriptional signatures were enriched in translation- and methylation-related processes, as well as WD40-domain-containing proteins. Interestingly, lipid catabolic pathways were preferentially augmented at early time points, whereas PI3K signalling and translational programs became more prominent at later stages, suggesting metabolic reprogramming associated with differentiation.

Overall, our findings highlight an early window of cellular plasticity after injury characterized by transient proliferation, followed by progressive activation of neuronal and glial differentiation programs during regeneration.

### CG14309 encodes an extracellular Heparanase-like protein early activated after injury

We hypothesized that early transcriptional changes in proliferative cells are associated with quiescence exit and the activation of regenerative programs. To test this, we focused on a subset of early injury-induced genes (**Figure 2A**) and assessed their contribution to the proliferative response by quantifying signal for the mitotic marker pH3. Gene knock-down (RNAi) was conditionally induced in adult flies by inactivation of thermosensitive (ts) Gal80ts, a Gal4 repressor. As expected, RNAi of *myc* during regeneration significantly reduced the number of PH3+ cells, as previously reported (Simões et al., 2022), which was also the case for knock-down of c*rumbs*, an inhibitor of the growth promoting Hippo pathway (Elbediwy et al., 2016) (**Figure 2B**). A similar effect was observed with RNAi of the fatty acid synthase *FASN2*, which has been linked to neural stem cell proliferation (Neumueller et al., 2011; Knobloch et al. 2013).

**Figure 2.**
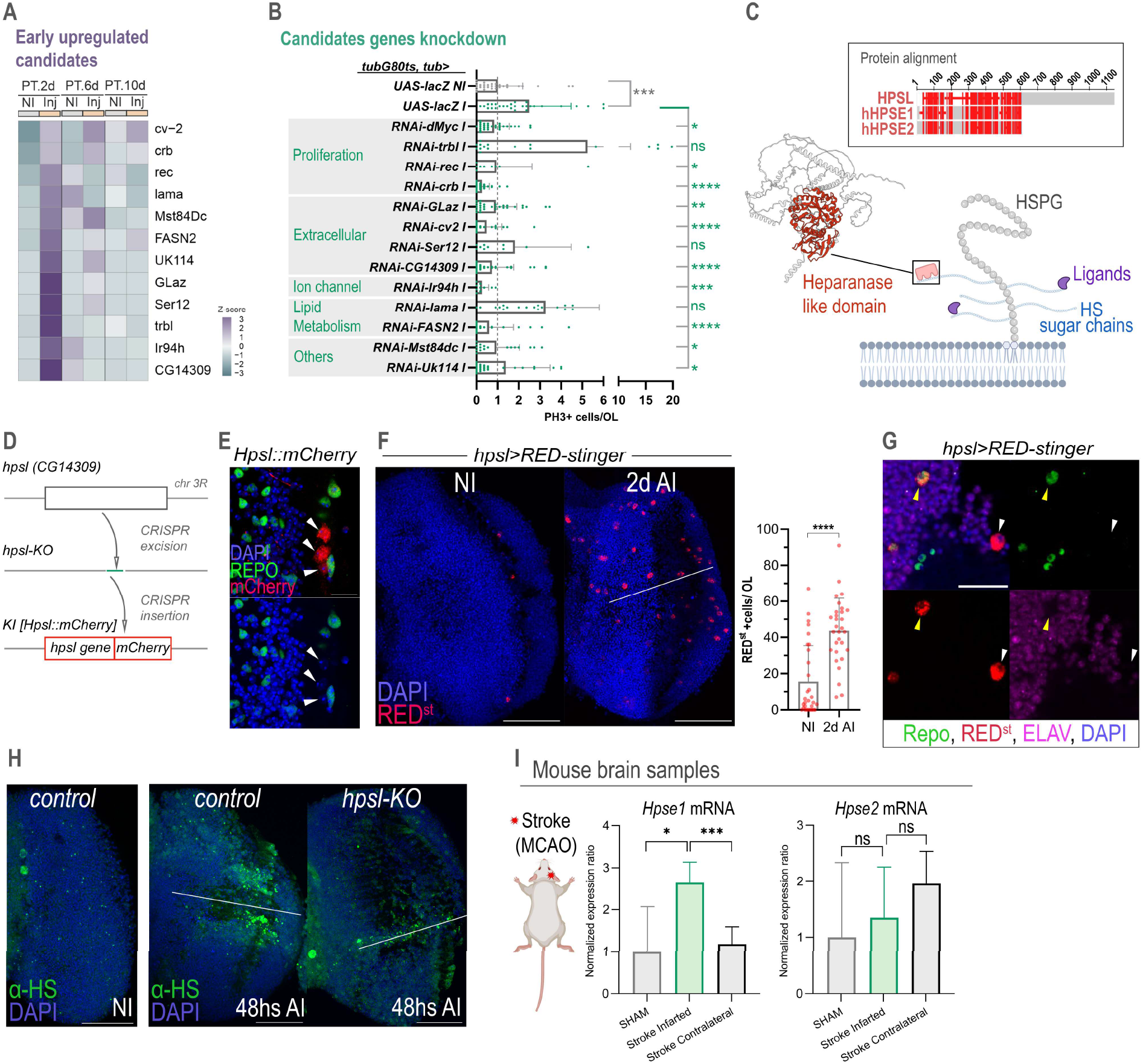
Identification of Hpsl as an injury-induced regulator of proliferation. (A) Heatmap showing early injury-upregulated candidate genes identified from transcriptomic analyses of proliferative cells. (B) RNAi-based functional screen of candidate genes. Quantification of pH3+ mitotic cells per optic lobe (OL) following knock-down of the indicated genes. Candidate genes were grouped according to predicted biological function. *Hpsl*/*CG14309* knock-down strongly reduced injury-induced proliferation. Data is shown as mean ± SD. 1-way ANOVA with Dunnet’s correction (vs *lacZ* injured animals) and p-values as follow: *p <0.05, **p <0.01, ***p <0.001, ****p <0.0001, non-significative (ns). (C) Structural analysis of Hpsl. Sequence alignment between *Drosophila* Hpsl and human heparanases (HPSE1 and HPSE2) reveals conservation within the heparanase-like domain. Schematic representation of Hpsl association with heparan sulfate proteoglycans (HSPGs). (D) Generation of *hpsl* mutant and reporter alleles using CRISPR-mediated genome engineering. (E) Representative image of the endogenous Hpsl::mCherry reporter in the adult OL. Repo (green) labels glial cells and DAPI labels nuclei. (F) Images of *hpsl*>RED-stinger expression in non-injured (NI) and injured (INJ) OLs. Graph: Quantification of RED+ cells per OL is shown on the right. Graph shows means and error bars represent standard deviation. T-test analysis and p-value ****p <0.0001. (G) Higher-magnification images of *hpsl*-expressing cells in injured (INJ) OLs. Repo, ELAV, and RED-stinger labeling identify the cellular distribution of *hpsl* expression. (H) Representative images of heparan sulfate (HS) staining in control and *hpsl* mutant OLs under no injury (NI) and injury (INJ) conditions. (I) Analysis of *Hpse1* and *Hpse2* mRNA expression in mouse brain hemispheres following transient middle cerebral artery occlusion (MCAO), causing ischemic injury. Expression levels were measured in sham-operated, infarcted, and contralateral brain tissue. Data is shown as mean ± SD. Welch’s t-test with Holm’s correction for multiple comparisons and p-value as follow: *p <0.05, ***p <0.001. Scale bars: 10µm (E), (G) and 50 µm (F) and (H).

Among the early activated factors, we further found genes coding for extracellular proteins including *glaz*, *cv2* and *CG14309* which were required for the proliferative response (**Figure 2B**). Remarkably, both Cv2 (mammalian Bmper) and CG14309 were predicted to interact with extracellular heparan sulfate proteoglycans (HSPGs), suggesting that ECM remodeling plays a critical role during the early repair response to brain injury.

Based on these findings, we further analyzed the properties of the uncharacterized gene *CG14309.* Sequence and protein folding analyses revealed a Heparan Sulfate (HS)-binding domain (AAs 20 to 425) homologous to mammalian Heparanases HPSE1 and HPSE2 (**Figure 2C**, **Figure S2A**), whereas the unstructured N-terminal part of CG14309 is not conserved and contains a potential furin cleaving site (AA 478) (**Figure S2A**). Thus, we hereafter refer to CG14309 as Heparanase-like (Hpsl).

Given the rapid induction of *hpsl* in early regeneration clusters, we further explored its function by generating *hpsl* knock-out (*KO*) flies by CRISPR-Cas9-mediated gene targeting, followed by knock-in (KI) of *hpsl-mcherry* (*KI[hpsl-mcherry]*) to capture expression under endogenous control elements (**Figure 2D**). Hpsl::mCherry protein was detected during late oogenesis (**Figure S2B**) consistent with reported transcriptional data sets (Leader et al., 2018). In the adult brain, we did not detect significant expression under homeostasis but observed injury-induced activation of Hpsl::mCherry at the lesion site (**Figure 2E**). We further used an *hpsl* enhancer trap line (*hpsl-GAL4*), which also showed transcriptional activation in late-stage oogenesis (**Figure S2B**) and was similarly upregulated acutely after brain injury, mainly in subsets of Repo+ glia and Repo- cells in the injured brain area (**Figure 2F**), whereas *hpsl* expression was never detected in Elav+ neurons (**Figure 2G**).

In mammals, HPSE1 promotes the cleavage of HS sugar chains attached to proteoglycans embedded in the extracellular space, whereas evolutionary more conserved HPSE2 retains high-affinity HS binding, but is catalytically inactive (Vahdatahar et al., 2025). To understand the braińs HS landscape upon which Hpsl may exert a modulatory role, we first attempted to map the location of HS sugar chains in the adult fly brain. *Drosophila* HS shows species-specific characteristics, including low O-sulfated domains (Kusche-Gullberg et al., 2012), but glycosylation patterns are difficult to survey *in vivo*. Nevertheless, *botv, a* key enzyme initiating the addition of HS glycosaminoglycan chains to core proteins, was widely expressed in intact and injured OLs, pointing to the presence of HS in the ECM (**Figure S2D**). Moreover, an antibody reported to recognize accessible sulfated HS-motifs appeared to detect increasingly exposed epitopes in brains of injured wild-type flies, which was not the case in lesioned *hpsl* KO animals (**Figure 2H**). Based on these results, we envisioned that Hpsl specifically interacts with HS sugar chains in the extracellular space following its rapid upregulation post injury.

We further investigated whether Heparanase activation is also detected in a mouse model of ischemic brain injury. Indeed, *Hpse1*, but not *Hpse2*, showed significant upregulation in the stroke-affected hemisphere compared to the contralateral side or sham controls (**Figure 2I**). In support of this, we also found increased *Hpse1* expression in brains of mice subjected to contusion and hemorrhagic stroke when analyzing a published single-cell dataset (Jha et al., 2024) (**Figure S2E**). Taken together, the findings show that induction of Heparanase-like factors acutely after brain injury is conserved across species, likely entailing alterations in extracellular heparan sulfate (HS) properties in the extracellular matrix (ECM).

### HPSL promotes tissue plasticity and animal recovery after brain injury

To further study the function of Hpsl in the extracellular space, we examined *hpsl* null flies, which are viable and show normal morphology and lifespan (**Figure S3A-B**), similar to *Hpse1* KO mice that have no obvious defects (Zcharia et al., 2009). However, flies devoid of Hpsl displayed increased vulnerability following brain injury, resulting in reduced survival compared to control flies (**Figures 3A and S3C**). When challenged with stab lesions, *hpsl* KO flies exhibited a significantly reduced proliferative response after injury, consistent with initial RNAi results (**Figure 3B and S3D**).

**Figure 3.**
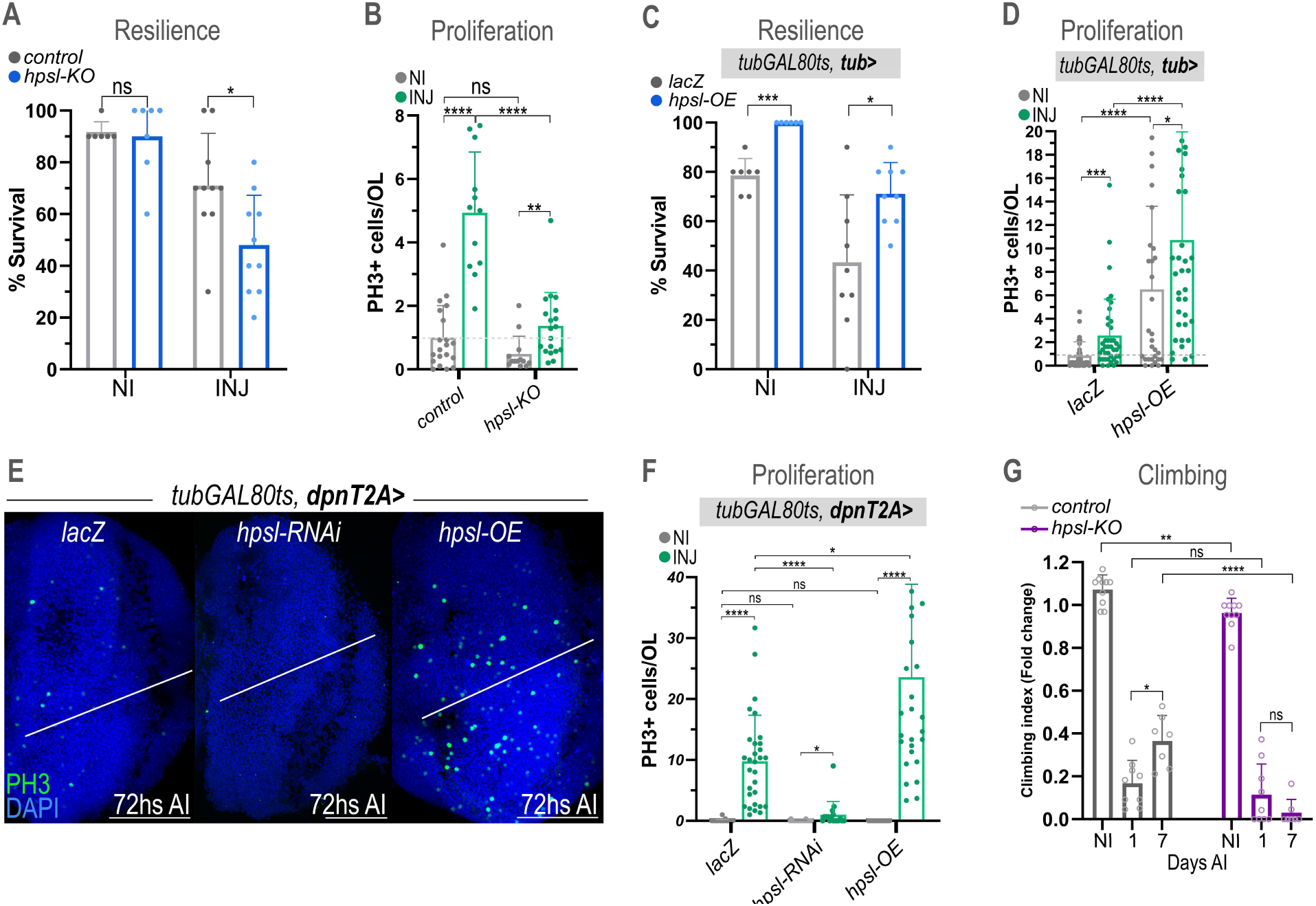
Hpsl promotes cell divisions, animal resilience and recovery after brain damage. (A) Survival analysis of control and *hpsl* KO animals under non-injured (NI) and injured (INJ) conditions. Means and standard deviation are shown. Every data point represents the mean of 1 vial with 10 flies. non-significant (ns) and *p <0.05 as determined by t-test analysis. (B) Quantification of pH3+ mitotic cells in control and *hpsl*-KO optic lobes (OLs) under NI and INJ conditions. Data is shown as mean ± SD. **** p <0.0001 and non-significant (ns) as determined by 2-way ANOVA with Dunn’s multiple comparison correction. (C) Survival analysis of animals overexpressing lacZ (control) and *hpsl* (*hpsl*-OE) in the indicated genotype (*tubGAL80ts, tub>*) under NI and INJ conditions. Every data point represents the average of 1 vial with 10 flies * p <0.05 and *** p <0.001, as determined by t-test. (D) Quantification of pH3+ mitotic cells in control (*lacZ*) and *hpsl-OE* animals under NI and INJ conditions. Graph shows means and error bars represent standard deviation. Kruskal–Wallis ANOVA with Dunn’s multiple comparison correction and p-value as follow: *p <0.05, **p <0.01, ***p <0.001, ****p <0.0001, non-significant (ns). (E) Representative images of adult OLs expressing control (*lacZ*), *hpsl-RNAi* or *hpsl-OE* constructs in progenitors (*tubGAL80ts, dpnT2A>*). pH3+ cells are shown in green. Nuclei are labelled with DAPI. (F) Quantification of pH3+ cells in the genotypes shown in (E). Means and standard deviation are shown. Kruskal–Wallis test with Dunn’s multiple comparison correction: *p <0.05, **p <0.01, ***p <0.001, ****p <0.0001, non-significative (ns). (G) Climbing assay performed at the indicated time points after injury in control and *hpsl*-KO animals. * p <0.05, ** p <0.01, **** p <0.0001 and non-significant (ns) as determined by 2-way ANOVA. Scale bars: 50µm (E)

In contrast, flies provided with an approx. 3-fold increase in *hpsl* expression at adult stage (*tub>hpsl-OE*; *tub>Gal80ts*)(**Figure S3E**), showed enhanced resilience to brain injury (**Figures 3C and S3F**) and exhibited increased cell divisions, both in injured and intact OLs (**Figures 3D and S3G**). To determine the cell type–specific requirement of Hpsl, we targeted its expression in neural progenitors and glial cells at adult stage using *dpnT2A-Gal4* and *repo-GAL4* drivers, respectively. *Hpsl* knock-down in either cell population significantly reduced mitotic counts after injury, suggesting that Hpsl facilitates regeneration via a coordinated function in glia and neural progenitors (**Figures 3E and S3H**). Remarkably, increasing *hpsl* expression in either cell population alone was sufficient to enhance injury-induced proliferation, underlining the capability of Hpsl as a strong plasticity driver in the adult brain (**Figures 3E and S3H**).

Finally, we interrogated how Hpsl activity impacts functional recovery after brain injury using a climbing assay (negative geotaxis), widely used to measure neurodegeneration or neural circuit dysfunction (Barone and Bohmann, 2013; Madabattula et al. 2015). Control animals with bilateral OL injury exhibited a marked impairment in climbing performance acutely (1 day) AI, followed by a partial recovery of this behavior by 7 days AI (**Figure 3F**). In contrast, *hpsl* mutants failed to exhibit improvements at delayed time points after injury. Altogether, these results establish that Hpsl does not only promote tissue plasticity and resilience to brain damage but can significantly contribute to recovery of behavior after injury.

### Extracellular Glypicans are rapidly remodeled after brain injury

HS Glycosaminoglycan chains are present on different core proteins present in the extracellular space, forming HS proteoglycans (HSPGs). Similar to mammals, flies produce three different HSPG forms (secreted, GPI-anchored and transmembrane) that could act as potential sources of extracellular HS, for which important roles in cell-cell signalling have been described (Lin and Perrimon, 2000). Available single cell data (Özel et al., 2021; Lago-Baldaia et al., 2023) supported expression of the HSPG glypicans *dally* (high) and *dlp (*low), syndecan (intermediate) and *perlecan/trol* (intermediate), especially in glial populations **(Figure S4A)**.

We found that Dally was highly expressed along glial extensions of Repo+ cells, forming a dense glial meshwork in the OLs as revealed by a Dally::GFP fusion protein with reported expression in neuropils (Pogodalla et al., 2021) (**Figure 4A**). Although signal was mainly detected in Repo+ glia, expression was also present in a subset of Repo- cells (**Figure 4B**). OL injury strikingly reduced the normally widespread Dally levels in extensive areas of the injured OL (**Figure 4A,C inset 1**), resulting in dynamic changes in the Dally pattern up to 1 week post the insult. In addition, we observed a pronounced Dally::GFP accumulation at the lesion site (**Figure 4C inset 2**) pointing to differential glial reactivity in proximity of the wound. The glypican Dlp, on the other hand, showed a distinct response, with low expression in homeostasis and a sharp increase in Dlp protein in the injured brain area (**Figure 4D, 4E**). The observed differences and non-overlapping expression (**Figure 4F**), indicate non-redundant roles of the two glypicans in injury-driven remodeling of the local tissue environment.

**Figure 4.**
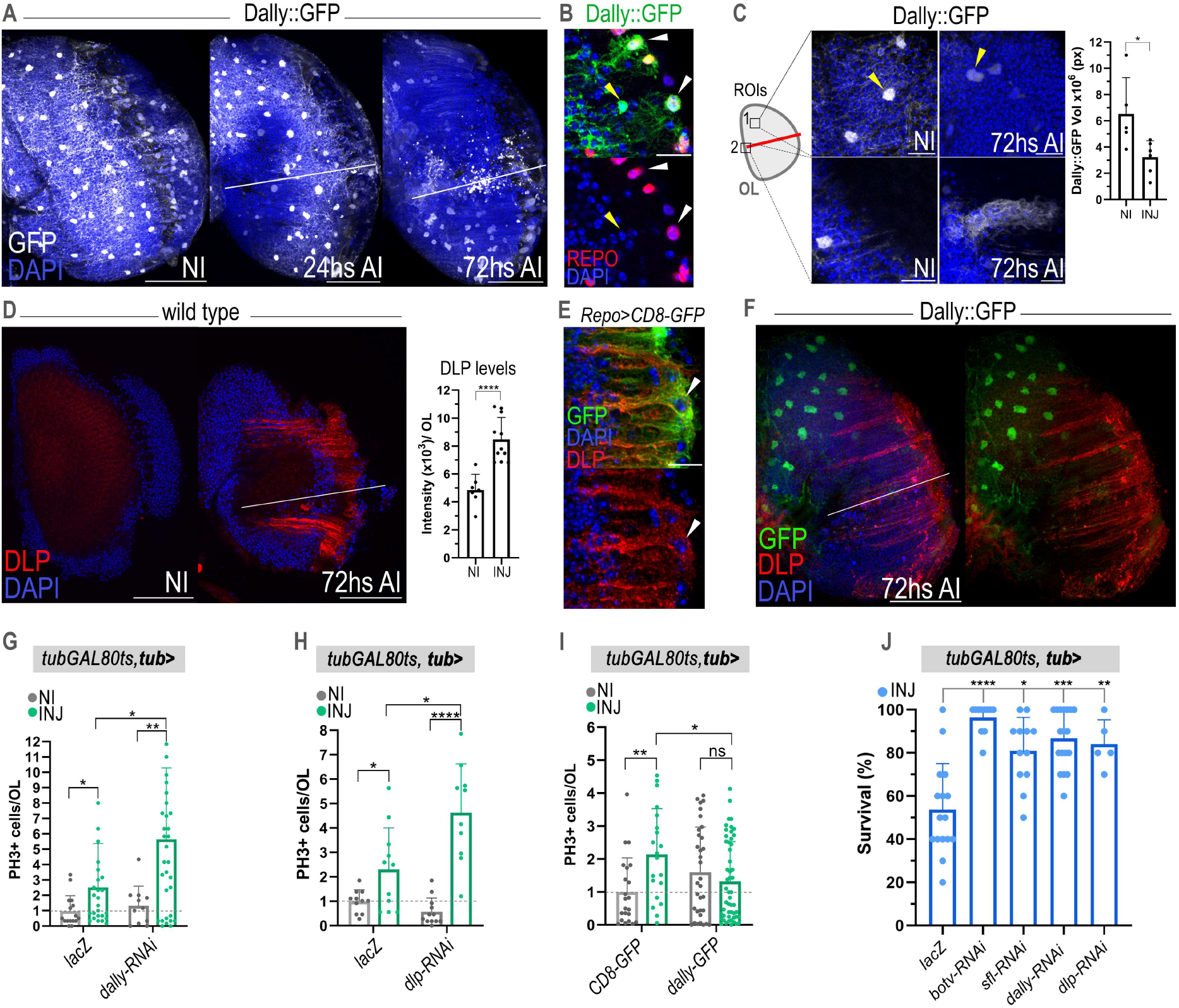
Injury-induced change in extracellular glypicans Dally and Dlp modulate the regenerative response in the adult brain. (A) Representative images of Dally::GFP expression in non-injured (NI) optic lobes (OLs) and at 24 h and 72 h after acute injury (AI). DAPI labels nuclei. (B) Higher-magnification images of Dally::GFP-positive cells in the injured OL. Repo labels glial cells. Repo+ (white arrowheads) and Repo- (yellow arrowheads) indicate representative Dally::GFP-expressing cells. (C) Representative regions of interest (ROIs) used to assess Dally::GFP expression in non-injured (NI) and injured (INJ) OLs. Arrowheads indicate representative Dally-positive cells. Quantification of the extent of Dally signal is shown on the right. Data is shown as mean ± SD. * p <0.05 as assessed by t-test (D) Representative images of Dlp immunostaining in non-injured (NI) and injured (INJ) OLs. Quantification of Dlp fluorescence intensity is shown on the right. Data is shown as mean ± SD. **** p <0.0001 as assessed by Mann-Whitney-Wilcoxon. (E) Representative image of *Repo>CD8-GFP* optic lobes stained for Dlp following injury. Arrowheads indicate Dlp-enriched regions associated with glial processes. (F) Representative image showing Dally::GFP expression (green) relative to Dlp accumulation (red) in the injured OL, DAPI stains nuclei. (G) Quantification of pH3+ mitotic cells per OL in control (*lacZ*) and *dally*-RNAi flies without injury (NI) and with injury (INJ). Data is shown as mean ± SD. Kruskal–Wallis test with Dunn’s multiple comparison correction. *p <0.05, **p <0.01. (H) Quantification of pH3+ mitotic cells per OL in control (*lacZ*) and *dlp*-RNAi flies without injury (NI) and with injury (INJ). * p <0.05; ** p <0.01, **** p <0.0001 as determined by Kruskal–Wallis with Dunn’s multiple comparison correction. (I) Quantification of pH3+ mitotic cells per OL in control (CD8-GFP) and overexpression of Dally-GFP flies under no injury (NI) and injury (INJ) conditions. * p <0.05; ** p <0.01 and non-significant (ns) as assessed by Kruskal–Wallis test with Dunn’s multiple comparison correction. (J) Survival rate of injured flies expressing the indicated RNAi constructs. Survival was monitored following acute injury and compared with control (*lacZ*) flies. ** p <0.01, *** p <0.001, **** p <0.0001 as assessed with 1-way ANOVA and Dunnet’s correction for multiple comparisons. Scale bars: 10 µm (B), (C), (E) and 50µm (A), (D) and (F)

To assess the causal role of glypican remodeling in the regenerative response, we manipulated *dally* and *dlp* expression in adult brains prior to injury, which in both cases resulted in increased injury-induced proliferation (**Figure 4G–H and S4B-C**). In contrast, activation of *tubulin*-driven *dally* overexpression in the adult brain restricted proliferative capacity after injury (**Figure 4H and S4D**), highlighting that broadly increased glypican levels crucially suppress injury-induced plasticity in the adult fly brain.

The plasticity-restricting effect of glypicans may arise from their long disaccharide chains that extend into the extracellular space, where they bind arrays of diverse ligands including growth factors, lipoproteins and ECM-regulators according to recent “HS-interactome” mapping (Gómez et al., 2021). In support of a strong contribution of the HS chains, we observed that adult onset targeting of glypicans (core protein + HS chains) similarly increased resilience to brain injury as conditional knock-down of HS initiating and modifying enzymes (Botv, Sfl) (**Figures 4J and S4E,S4F**), whereas no effect was detected on survival in unchallenged conditions (**Figure S4G**).

Together, these results demonstrate that dynamic remodeling of Heparan-sulfate carrying glypicans critically regulate the injury-adaptive response of brain tissue by enabling proliferative capacity and responses that contribute to organismal fitness after brain insult.

### Hpsl locally regulates critical Dpp/BMP signalling to drive regeneration

Glypicans and other HSPGs have been shown to act as storage site for growth factors or serve as co-receptors to facilitate signalling by FGF, Wnts and BMP/TGF-Beta family ligands (Lin and Perrimon, 2000).

In *Drosophila*, the Glypican Dally has been identified as a key extracellular regulator of signalling by Dpp, the fly orthologue of mammalian TGF-β/BMP ligands (Jackson et al., 1997; Belenkaya et al., 2004; Affolter and Basler, 2007; Vuilleumier et al., 2010; Hamaratoglu et al., 2014). Given the upregulation of Dpp pathway components *(e.g. cv-2, pent)* in early regenerative clusters **(Table 1)**, we hypothesized that ECM modifications prompted by Hpsl and glypican remodeling may alter Dpp/BMP pathway activity in the injured brain (**Figure 5A**). Indeed, we observed consistent injury-dependent activation of the Dpp-pMad downstream target *dad*-GFP, reporting Dpp/BMP signalling (Hamaratoglu et al., 2011) (**Figure 5B**). We detected activation of the reporter predominantly in glial cells (∼90% Repo+) and also in a few Repo- cells (**Figure 5C**). As expected, injury-induced Dpp/BMP pathway activity was impaired upon *dpp* RNAi or overexpression of the Dpp pathways inhibitor Brinker (**Figures 5D and S5A**), confirming Dpp-mediated dad-GFP activation upon injury.

**Figure 5.**
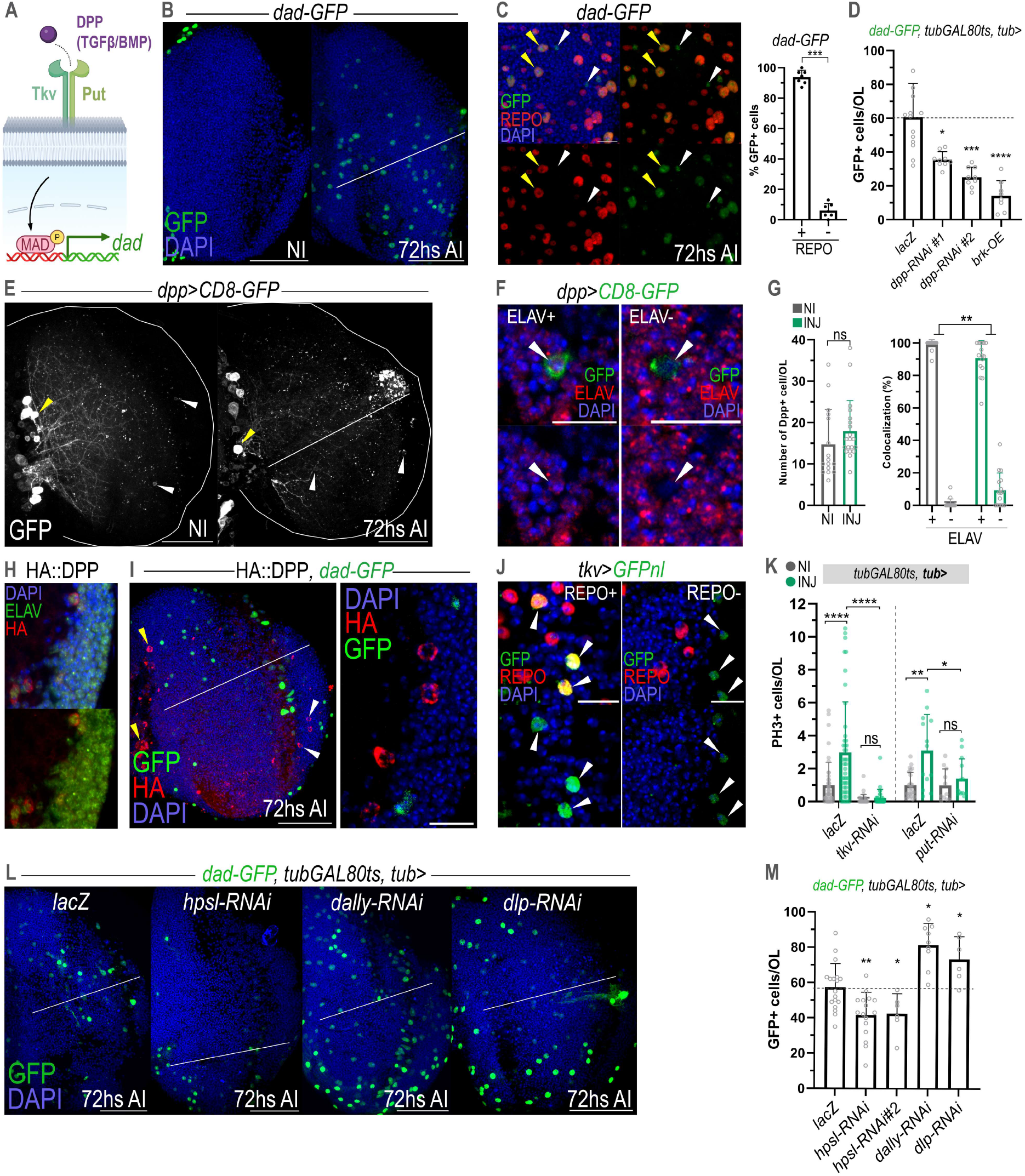
Dpp/BMP signalling is activated after injury and promotes regeneration through HPSL and Dally-dependent regulatory mechanisms. (A) Scheme of the *Drosophila* BMP/TGF-β signalling pathway. Binding of Dpp to the receptors Thickveins (Tkv) and Punt (Put) promotes Mad phosphorylation and activation of downstream target genes, including *dad*. (B) Representative images of the dad-GFP reporter in non-injured (NI) and injured optic lobes (OLs) 72 h after injury (AI). (C) Higher-magnification of dad-GFP+ cells in injured OLs. Repo staining is shown in red and DAPI in blue. Repo+ (yellow arrowheads) and Repo- (white arrowheads) indicate representative reporter-positive cells. To the right, quantification of the percentage of dad-GFP+ cells that colocalize with Repo per OL after injury. (D) Quantification of the number of dad-GFP+ cells per OL after injury in the indicated genotypes. Data is shown as mean ± SD. ** p <0.01, **** p <0.0001 as assessed by One-way ANOVA with Dunnett’s multiple comparisons (E) Representative images of *dpp>CD8-GFP* expression in non-injured OLs (NI) and 72hs after injury (AI). Yellow arrowheads indicate GFP+ cell bodies from pacemaker lateral neurons. White arrowheads point to cells within the OL. (F) Higher-magnification images showing the colocalization between *dpp>CD8-GFP*-positive cells and the neuronal marker ELAV. Arrowheads indicate representative GFP+ cells within the OL with positive and negative ELAV mark. (G) Quantification of *dpp>CD8-GFP*-positive cells per OL and their colocalization with ELAV without injury (NI) and 72hs after injury (INJ). Data is shown as mean ± SD. non-significant (ns) (graph 1) as assessed by Mann–Whitney–Wilcoxon and ** p <0.01 (graph 2). (H) Neurons with HA::DPP expression (red) in the adult OL. ELAV expression is shown in green. Cell nuclei are counterstained with DAPI (blue). (I) Representative images of HA::DPP (red) and dad-GFP expression 72 h after injury (AI). Right panel shows a higher magnification. Arrowheads indicate representative HA::DPP-positive cells. (J) Representative images of *tkv>GFPnl* reporter expression in Repo+ and Repo- cells. Arrowheads indicate representative GFP+ cells. (K) Quantification of pH3+ mitotic cells per OL following knock-down of BMP signalling components in non-injured (NI) and injured flies (INJ) (72h AI). Data is shown as mean ± SD. * p <0.05; **** p <0.0001 as assessed by Kruskal–Wallis test with Dunn’s multiple comparison correction. (L) Representative images of dad-GFP reporter expression following knockdown of *hpsl*, *dally*, or *dlp*. Nuclei are counterstained with DAPI (blue). (M) Quantification of dad-GFP+ cells per OL in the indicated genotypes. Data is shown as mean ± SD. * p <0.05; ** p <0.01, *** p <0.001 as assessed by One-way ANOVA with Dunnett’s multiple comparisons. Scale bars: 10µm (C), (F), (H), (J) and 50µm (B), (E), (I) and (L)

To trace the cellular source of Dpp in the adult brain, we analyzed dpp-driven membrane GFP (*dpp>CD8-GFP*). We observed previously described Dpp-producing pacemaker neurons, lateral to the central brain (Polcowñuk et al., 2021) (**Figures 5E and S5B**), which extend prominent projections into the OLs, suggesting Dpp delivery under physiologic conditions. In addition, we detected a small subset of normally active Dpp-producing neurons within the OLs (**Figure 5F**). Interestingly, Dpp-expression was not induced in further neurons (Elav+) upon injury, although very few GFP+ Elav- cells were detected after damage (**Figure 5G**). In all conditions, Dpp signal never overlapped with Repo+ glia (**Figure S5C**). Similarly, endogenous HA-tagged Dpp (HA::Dpp) (Matsuda et al., 2021) was strongly enriched in pacemaker lateral neurons and a restricted set of OL neurons in the medulla, the numbers of which did not change after injury (**Figures 5H and S5D**). The combined visualization of HA::Dpp ligand-producing neurons and Dpp pathway activation (dad-GFP) predominantly in glia (**Figure 5I**), strongly suggested a non-cell autonomous role of Dpp signalling in response to brain damage. This was consistent with the expression of the main Dpp receptor (Tkv) mainly, but not exclusively, in glial cells, in intact and lesioned OLs (**Figure 5J and S5E-F**). Importantly, blocking of Dpp signalling by conditional RNAi of the BMP type I receptor *thickvein (tkv)* (Ruberte et al., 1995) and the type II receptor *punt* (*put*) reduced the number of observed cell divisions at the lesion site (**Figure 5K**). Taken together, the results establish that injury leads to Dpp-mediated neuro-glial interactions, able to drive a proliferative response in the normally non-permissive adult brain environment.

Although Dpp and its receptors are constitutively present in the adult brain, pathway activation is only observed following injury, hinting at a molecular mechanism that promotes ligand mobilization and redistribution from its original sources to responsive target cells. Based on our results, we considered Hpsl binding to HS chains as an important event that could rapidly liberate Dpp from HS storage sites in the ECM. To formally test this, we targeted *hpsl* with two RNAi lines and monitored the effect on Dpp signalling. Indeed, Hpsl inhibition significantly compromised the engagement of the Dpp pathway in injured brains, showing that Hpsl crucially facilitates Dpp-dependent cellular cross-talk (**Figure 5L-M**). Interestingly, conditional RNAi of *dally* or *dlp* also enhanced Dpp/BMP signalling (**Figure 5L-M**), potentially due to reduced extracellular Dpp sequestering, which is in line with the observed increase in proliferation following injury (**Figure4G-H**). The findings therefore show that injury-driven alterations in ECM properties lead to rapid engagement of pro-regenerative Dpp signalling across neuro-glial circuits, likely due to increased availability and/or diffusion of limited Dpp growth factor in the extracellular space.

### HPSL promotes progression of invasive brain tumors

Tissue repair responses are frequently co-opted by cancer cells as both regeneration and tumorigenesis rely on common gene regulatory networks to sustain proliferation, nutrient availability and ECM remodeling (Wong and Whited, 2020; Floc’hlay S et al., 2023; MacCarthy-Morrogh and Martin, 2020). Accordingly, perduring activation of an “injury response program” has recently been detailed in glioblastoma (Hamed et al., 2025).

To investigate whether Hspl is recruited by cancer cells, we suppressed its function in an aggressive *Drosophila* glioblastoma (GB) model, in which constitutively active insulin and EGFR axes in glia (*repo>EGFRλ, PI3KCAAX*) drive massive glial proliferation and infiltration by glial microtubule-like protrusions, ultimately inducing neuronal apoptosis (Read et al. 2009; Witte et al., 2009) (**Figure 6A**). Strikingly, targeting of *hpsl* in glia significantly restrained GB progression, leading to lower GB cells counts in larval brains (**Figure 6A**). Suppression of HPSL function also reduced the growth of GB when activated in adult flies, in the far less growth-permissive environment of the mature brain, as assessed 15 days after GB induction (**Figures 6B and S6A**). *Hpsl* RNAi also markedly inhibited glial overgrowth driven by the oncogenic version of Ras (*rasV12*) (**Figures S6B-C**). Importantly, glioblastoma-bearing flies exhibited impaired climbing ability 10 days after tumor activation (**Figure 6C**) and premature lethality (**Figure 6D**), both of which were significantly reduced upon *hpsl* silencing. Altogether, these findings show that Hpsl promotes plasticity and glioblastoma progression and suggest shared roles of ECM-driven tissue plasticity upon injury and in the tumor microenvironment.

**Figure 6.**
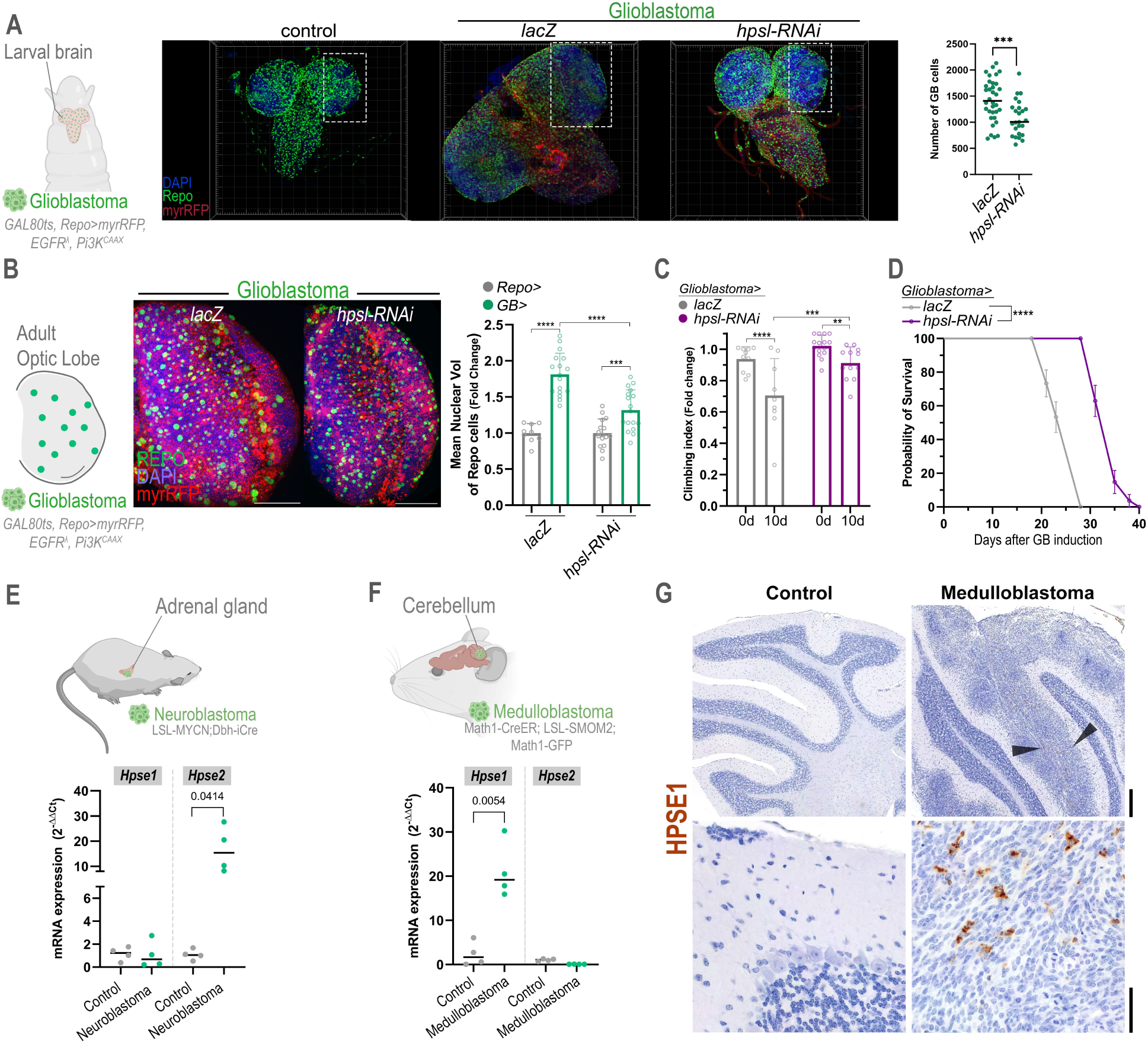
Hpsl-dependent mechanisms regulate tumor growth in the nervous system. (A) Representative images of larval brains from control animals and a *Drosophila* glioblastoma (GB) model expressing activated EGFRλ and PI3KCAAX in glial cells (repo>) with *lacZ* and *hpsl* knockdown. Quantification of glioblastoma cells is shown on the right. ***p <0.001 as assessed by t-test. (B) Representative images of adult-induced optic lobe glioblastoma (*Gal80ts; repo>myrRF, EGFRλ,PI3KCAAX*) in control and *hpsl*-RNAi animals. Quantification of the nuclear volume of Repo+ cells is shown on the right. Data is shown as mean ± SD. 2-way ANOVA and p-value as follow: ***p <0.001, ****p <0.0001. (C) Climbing assay performed at the indicated time points following glioblastoma induction in *lacZ* and *hpsl*-RNAi animals. **p <0.01, ***p <0.001, ****p <0.0001 as assessed by 2-way ANOVA. (D) Survival curves of glioblastoma-bearing flies expressing *lacZ* or *hpsl-RNAi* constructs. ****p< 0,0001 as determined by Mantel-Cox test. (E) Analysis of *Hpse1* and *Hpse2* mRNA expression in a mouse neuroblastoma model (LSL-MYCN;Dbh-iCre ) compared with control tissue (n= 4 for both conditions). Welch’s t-test and p-value is indicated. (F) Analysis of *Hpse1* and *Hpse2* mRNA expression in a mouse medulloblastoma model (Math1-CreER; LSL-SMOM2; Math1-GFP) compared with control cerebellar tissue (n= 4 for both conditions). Welch’s t-test and p-value is indicated. (G) Representative microphotographs of normal mouse cerebellum and medulloblastoma-bearing cerebellum (n = 6 medulloblastoma; n = 2 controls). Arrowheads indicate tumor areas with positive HPSE1/heparanase immunoreactivity. Lower panels show higher-magnification views of HPSE1 immunostaining, highlighting absent signal in control cerebellum and multifocal positivity in medulloblastoma. DAB counterstained with hematoxylin. Scale bars: 50µm (B), 200 µm (G upper panel) and 40 µm (G bottom panels)

To determine whether members of the Heparanase family are similarly upregulated in invasive nervous system tumors in mice, we performed quantitative expression analyses comparing aggressive neuroblastoma samples generated by Cre-conditional activation of MYCN in dopamine β-hydroxylase-expressing cells (LSL-MYCN;Dbh-iCre) (Althoff et al., 2015) with normal adrenal gland tissue, as well as, sonic hedgehog medulloblastoma (SHH MB) samples from Math1-CreER; LSL-SMOM2; Math1-GFP mice (Schüller et al., 2008; Tiberi et al., 2014) with normal cerebellum. Neuroblastoma cells displayed increased *Hpse2* expression without significant changes in *Hpse1* levels (**Figure 6E**). In contrast, the medulloblastoma (MB) showed strong upregulation of *Hpse1* (∼20-fold), whereas *Hpse2* expression remained unchanged (**Figure 6F**). We then analyzed HPSE1 protein expression *in vivo*, in brains of age-matched animals, at the time when motor symptoms appear (5 weeks after tumor initiation) in the pediatric medulloblastoma model. HPSE1 was specifically and consistently increased in cells at the midline of the dense and proliferative tumor mass (**Figure 6G**), with no appreciable expression in control cerebellum samples (n = 6 MB; n = 2, controls).

Together, these results indicate that Heparanase-driven programs form part of the advanced tumor signature, which are associated with malignant NS invasion and reminiscent of regenerative responses following tissue damage.

## Discussion

The restricted regenerative capacity of the adult brain remains a major barrier to functional recovery after injury, yet which components of the regenerative tissue environment regulate plasticity is largely unknown. By combining transcriptional fingerprinting of dividing cells with functional genetics, we uncovered an ECM-based plasticity-conferring mechanism in the injured fly brain. We show that rapid release of the Heparan Sulfate (HS)-interacting factor Hpsl **(Figures 2 and 3)** and dynamic modification of the extracellular glypican landscape (**Figures 4)** enable the harnessing of limited Dpp/BMP growth factor for neuro-glial cross-talk, which promotes activation of latent, proliferation-competent cells **(Figure 5)**. Conversely, lack of Hpsl function or widespread Dally/glypican overexpression significantly restrict injury-induced plasticity in the injured brain area and reduce organismal resilience to brain injury and recovery of behavioral control.

Our findings support a model in which injury-induced Hpsl binds extracellular HS chains to release sequestered Dpp/BMP ligands. Together with glypican remodeling, including Dally downregulation, this can facilitate ligand diffusion and activation of Dpp/BMP signalling in glia and neural progenitors to drive proliferation (**Figure 7**).

**Figure 7.**
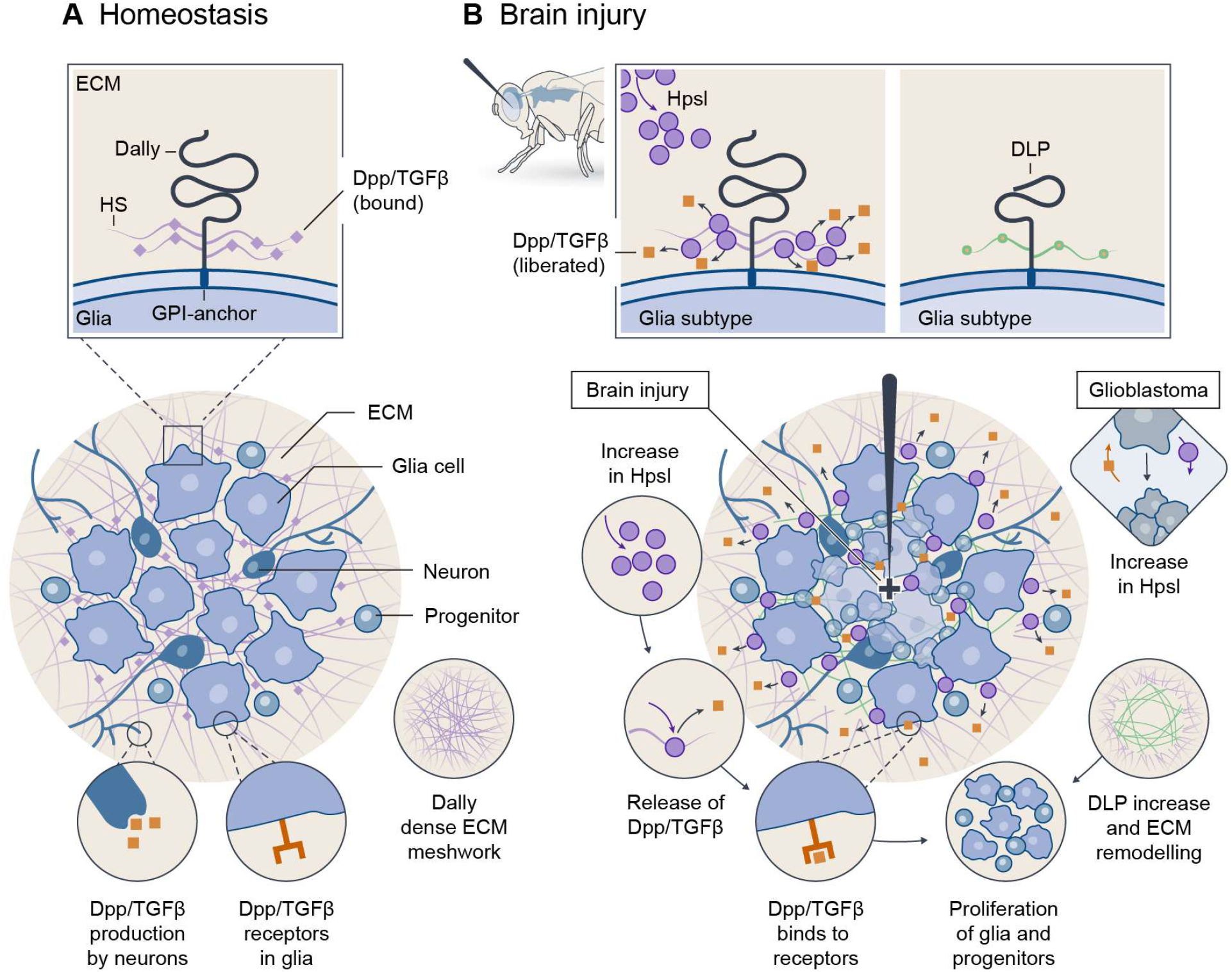
Hpsl and Glypican-related ECM remodelling controls brain plasticity. (A) Under homeostatic conditions, the extracellular space is characterized by extensive expression of the Heparan-Sulfate (HS)-decorated glypican Dally in glial networks. Dpp production and embedding in the ECM occurs via a restricted set of neurons. Although the Dpp/BMP receptor is widely expressed in glial cells (left lower corner), receptor engagement and Dpp/BMP signalling is not activated. (B) Upon brain injury, Hpsl is induced and interacts with extracellular HS. Acute damage leads to broad downregulation of Dally, whereas the glypican Dlp is upregulated in the injured brain area. Hpsl action and HS-related ECM remodeling lead to Dpp/BMP mobilization and activation of pro-regenerative Dpp/BMP signaling in glia and neural progenitors, triggering their proliferation. Hpsl function also facilitates the expansion of tumor cells in invasive glioblastoma (right corner).

However, additional signalling and biomechanical effects of Hpsl expression and ECM remodelling are likely to contribute to the beneficial effects on brain resilience and recovery from brain injury. Apart from the generation of new cells, the rewired extracellular space may additionally favor synaptic plasticity post injury. Such effects could be mediated via recently described effects of astrocyte-derived glypicans that actively stabilize synapses in neural circuits in the adult mouse brain (Bosworth et al., 2025) or HS-dependent functions of Neurexins in synapse organization (Zhang et al., 2018).

In flies, glypican HSPGs have been shown to regulate ligand distribution, availability and receptor engagement within the extracellular space of epithelia (Restrepo et al., 2014 ; Yan and Lin, 2009), underscoring their potential to modulate signalling. Intriguingly, we did not observe upregulation of Dpp/BMP or activation in additional cellular sources in response to brain injury **(Figure 5).** Instead, regulation occurred primarily at the level of extracellular components. Hspl function thereby provides the cell with the possibility to rapidly alter growth factor accessibility, which precludes the need for their *de novo* synthesis, thereby allowing a timely response to acute injury. Our findings further emphasize that the biological activity of BMP ligands cannot be solely inferred from expression patterns. Instead, BMP signalling output is critically shaped by ligand distribution and diffusion range, which are themselves determined by the properties of the extracellular space (Tønnesen et al., 2023) and its effects on the activity of BMP signalling components (Umulis et al., 2009; Restrepo et al., 2014; Sedlmeier and Sleeman, 2017).

The observed differential regulation of the two glypicans *dally* and *dlp* is intriguing (**Figure 4**). Although similar in structure, their affinity for growth factors may differ, as well as their level of HS-dependency (Nakato et al., 2024). Dally has been strongly implicated in Dpp signalling (Restrepo et al., 2014, Wartlick et al., 2011), whereas Dlp is more commonly associated with Wnt pathway modulation, at least in fly epithelia (Waghmare et al., 2020; McGough et al., 2020). Nevertheless, the fact that either knock-down of Dally or Dlp facilitated proliferation suggests that both may sequester growth factors, although they may function in a cell-type-specific manner, which remains to be further investigated.

Interestingly, Hpse1 expression has also been reported in proliferating cells in the peri-infarct region of the mouse brain early after stroke (Li et al., 2012), similar to what we found in this study **(Figure 2).** However, the functional consequences remained largely unexplored.

Our results suggest that Hpsl and the glypican Dally can synergistically promote Dpp/BMP pro-regenerative signalling and support brain repair. BMP pathway activity has also been found to regulate neural stem cell behavior and brain plasticity. However, under physiologic conditions, BMP pathway activity mainly regulates stem cell quiescence (Lim et al. 2000; Mira et al. 2010; Colak et al., 2008) suggesting that BMP signals provided by the neurogenic niche may activate different target genes.

In the context of brain cancer development, we found that Hpsl function is strongly linked to glioblastoma progression in fly models, promoting proliferation and tumor expansion that lead to behavioral decline and premature death. A conserved function of HPSEs in the mammalian brain is supported by their specific upregulation in invasive nervous system cancers, in both of our generated mouse models. HPSE1 has been previously associated with tumor progression including glioblastoma and medulloblastoma (Xiong et al., 2020; Zetser et al., 2003), with high human *Hpse 1* expression in primary GB correlating with low median survival (36.9 months) compared to increased median survial (98 months) of patients with low *Hpse1* levels (Bowman et al., 2017). Nevertheless, functional experiments to inhibit Heparanase in the brain tumor microenvironment have not been performed, although treatments of pediatric tumor cells with Heparanase inhibitors in vitro showed promising effect in reducing proliferative and invasive capacity (Spyrou et al. 2017). On the other hand, the proliferation-promoting role of the BMP/TGF-Beta pathway is widely recognized (Massague, 2008) and BMP inhibitors are tested in clinical trials for brain cancer as they may prove effective in targeting glioma-initiating cells (Anido et al. 2010).

Restricting the invasiveness of GB is highly relevant as these aggressive tumor cells infiltrate healthy brain tissue, which is the major reason they remain fatal despite resection. On the other hand, progression to metastatic behavior is associated with lower survival of children with medulloblastoma (Xiong et al., 2020). In this regard, the *hpsl* knock-down experiments in the *Drosophila* GB model provide initial proof-of-principle results for beneficial effects of Heparanase inhibition in invasive brain cancers, which encourage further testing in mouse models.

Together, our findings uncover a conserved ECM-based mechanism that coordinates regenerative signalling after brain injury, providing a strong framework for enhancing brain repair and limiting the regenerative programs co-opted by brain tumors.

Given the conserved nature of the components, the injury-induced Hpsl–Dally–Dpp/BMP axis may represent a conserved regulatory module of brain plasticity that can be harnessed to increase brain restorative processes or curb brain tumors.

## Acknowledgments

We are grateful to Henrique Veiga-Fernandes and David Bréa for mouse brain tissue. We thank the CF Molecular Tools Platform for support with cloning and transgenesis and the CF Imaging and Fly platform for technical support. We thank Markus Affolter, Christian Klaembt and Fisun Hamaratoglu for providing important fly lines, the DSHB Antibodies and Bloomington and Vienna *Drosophila* stock centers for reagents and fly lines. This work has been supported by grant HR23-00860 from LaCaixa & Fundação para a Ciência e Tecnologia (FCT), the ERC-Portugal program (FCT) to C.R. and the Champalimaud Foundation and PID2022-139786OB-I00 (MICINN) to S. C. This study was further supported by the Human Frontier Science Program (HFSP) Fellowship LT0014/2024-L (J.A.S.) (DOI:10.52044/HFSP.LT00142024), and FCT PhD fellowship to M.S. (PD/BD/114048/2015). A.S.D, SMF and PDNB are supported by QuantOCancer Project Horizon European Union’s Horizon 2020 programme (grant agreement No 810653). S.M.F. is supported by FCT (grant agreement 2025.02089.BD). This project was also supported by a FCT grant (2022.09083.PTDC) and AstraZeneca Award in Oncology and Liga Portuguesa Contra O Cancro to A.S.D.

## Author contributions

This study was conceptualized by C.R. with contributions from J.A.S., S.C. and A.S.D. Investigation and formal analyses were performed by J.A.S., M.S., A.C.O., D.A., S. F.M., P.D.N.B., T. C. and S.C. The manuscript was written by C.R. and J.A.S. with contributions from A.S.D. and S.C.

## Disclosure and competing interest statement

The authors declare no competing interests.

## Material and Methods

### Fly strains and husbandry

Flies (Drosophila melanogaster) were raised on standard cornmeal agar medium and maintained in an incubator set at 25 °C, 60% humidity with a 12h light/12h dark cycle or at 18°C when combined with *tub-Gal80ts*.

The following *Drosophila* lines were used:

Knock-down lines (*UAS-RNAi*): *UAS-cdk1* (BL 28368), *UAS-myc-RNAi* (VDRC 71133), *UAS-trbl-RNAi* (BL 42523), *UAS-rec-RNAi* (v106671), *UAS-crb-RNAi* (v39177), *UAS-GlaZ-RNAi* (BL 67228), *UAS-Ser12-RNAi* (BL 41624), *UAS-cv2-RNAi* (v109915), *UAS-CG14309-RNAi* (BL 67855) (also mentioned as UAS-hpsl-RNAi), *UAS-Ir94h-RNAi* (v100407), *UAS-lama-RNAi* (v107629), *UAS-FASN2-RNAi* (v105855), *UAS-Mst84dc-RNAi* (v23559), *UAS-UK114-RNAi* (v106775), *UAS-dally-RNAi* (BL 28747), *UAS-dlp-RNAi* (BL 50540), *UAS-botv-RNAi* (BL 61257), *UAS-sfl-RNAi* (BL 50538), *UAS-dpp-RNAi* #1 (VDRC 330518), *UAS-dpp-RNAi* #2 (BL 95276), *UAS-tkv-RNAi* (BL 40937), *UAS-put-RNAi* (39028), *UAS-CG14309*-RNAi (VDRC 103061) (mentioned as *UAS-hpsl-RNAi* #2).

Overexpression lines (UAS): *UAS-lacZ*, *UAS-RASV12, UAS-CD8-GFP, UAS-brk-OE* (gifts from Eduardo Moreno), *UAS-CD2-RFP* (BL 8529) *tub-Gal80ts (*BL7019), *UAS-RedStinger* (BL 8546), *UAS*-*Dally-GFP* (BL 94569), *UAS-GFP.nls* (BL 4776), *UAS-hpsl-OE* (BL 31839), *UAS- Pi3kCAAX*, *UAS-EGFR-λ* (gift from Sergio Casas-Tintó), Driver lines (Gal4) *hpsl-Gal4* (Kyoto 104733), *dpnT2A-Gal4* (Rhiner lab), *repo-Gal4 (*BL 7415) , *tub-Gal4*, *dpp-GAL4* (BL 93385), *tkv-GAL4* (BL 63958), *botv-GAL4* (BL 80617),

Reporter lines Source: *E(spl)mBetaHLH-GFP* (BL 65294), *KI[hpsl::mCherry]* (generated in this study), dad-GFP nl (nuclear), gift from Fisun Hamaratoglu, Dally::GFP (gift from Christian Klaembt), Dpp::HA (gift of Markus Affolter).

Other lines: Canton S (gift from Eduardo Moreno), *hpsl-KO* (generated in this study). Perma-Twin flies: *w; FRT40A, UAS-CD8-GFP, UAS-CD2-mir/ CyO; actinGal4, UAS-flp/TM6B* and *w; FRT40A, UAS-CD2-RFP, UASGFP-mir/ CyO; tubGal80ts/TM6B*.

### Mouse lines and maintenance

Mouse colonies were maintained in a certified animal facility in accordance with European guidelines for the laboratory animal use and care based on the 2010/63/EU Directive. Experiments involving mice presented in this work were approved by the Animal Welfare and Ethics Body and Direção-Geral da Alimentação e Veterinária (DGAV,Portuguese Authority).

### Generation of hpsl KO and KI flies

To create *CG14309/hpsl* KO flies (ChrIII) by CRISPR/Cas9-mediated gene editing, 1 kb 5’ and 3’ homology arms (HA) for *hpsl* were amplified from nos-Cas9 ChrII fly’s genomic DNA and directionally and sequentially cloned into pTV3 plasmid; HA’s plus the attP landing site and mCherry transgenesis marker were flanked by the Cas9 sgRNA target sequence for linearization of the Donor fragment, the template for Homology-Directed Repair. The two guide RNAs (sgRNA; sgRNA ATG-565, and sgRNA STOP-59) were cloned into the dual gRNA vector pCFD5 (Addgene #73914) by means of a ligation-independent cloning. All positive clones were identified by their restriction profile and sequenced. DNA was injected into nos-Cas9 (ChrII) flies.

For the *hpsl* knock-in line (Hpsl::mCherry), we created a white+ selection marker reintegration vector for *hpsl*, which restored the 565bp upstream of the *hpsl* ATG plus *hpsl* CDS+introns (3050bp) deleted in the KO, and insert mCherry in frame before the *hpsl* stop codon. *hpsl* was amplified from nos-Cas9 ChrII gDNA. The final Hpsl::mCherry KI sequences was assembled in two steps. 1. Gibson Assembly was used to fuse a 3’ portion of *hpsl* and mCherry and directionally clone them into the RIV-White vector between NheI and KpnI sites. 2. 5’UTR-partial *hpsl* was directionally cloned into the previous vector, between EcoRI and NheI. Positive clones were identified by their restriction profile and sequencing and injections performed into the *hpsl-*KO fly stock.

### Stab lesions

A thin sterile filament (0,1mm Fine Science Tools), wiped with 70% ethanol was introduced through the right eye (unilateral) or both eyes (bilateral injury) of at least 3–4 days-old adult CO2- anesthetized flies to the level of the medulla in the optic lobe as previously described (Fernández- Hernández et al, 2013).

### MCAO model (stroke)

For the stroke model (middle cerebral artery occlusion; MCAO) (Jackman et al., 2011), 2-month-old C57BL/6J mice were subjected to transient focal cerebral ischemia using an intraluminal filament, and samples analyzed 72hs later. The brain samples were obtained and processed as previously described (Simões et al., 2022).

### Perma-twin clones

To generate the PT flies, males from *w; FRT40A, UAS-CD8-GFP, UAS-CD2-mir/ CyO; actinGal4, UAS-flp/TM6B* were crossed to virgin females of the *w; FRT40A, UAS-CD2-RFP, UASGFP-mir/ CyO; tubGal80ts/TM6B* (Fernández-Hernández et a., 2013). The crosses were maintained at 18°C during development. Progeny was collected in separate vials and kept at this temperature until the age needed. 3-5 days old flies were selected and shifted to 29°C to activate the system and injured on the following day.

### Immunohistochemistry and antibodies

Fly brains were dissected in chilled PBS, fixed in 4% PFA for 20 min at RT, washed with PBS-Triton-X 0.4% and incubated with primary (overnight at 4 °C) and secondary antibodies (overnight at 4 °C). DAPI (1:1000) was used for counterstaining. Adult brains were mounted with a spacer to avoid compression of the tissue. For pH3 staining, samples were fixed overnight at 4 °C in 2% PFA, washed and incubated for 48h- 72h with primary Ab. Samples were imaged on an 880 Zeiss Confocal microscope, and images were analyzed using ImageJ and Zen Digital Imaging for Light Microscopy.

The primary antibodies used were: rabbit phospho-histone H3 (Ser10) 9701s Cell Signaling (1:50); mouse anti-Repo 8D12 (DSHB) (1:20); mouse anti-Elav 9F9A9 (DSHB) (1:50); rat anti-Elav 7E8A10 (DSHB) (1:50); mouse anti-Dlp 13G8 (DSHB) (1:20) and mouse anti-HS F58-10E4 Abcam (1:100).

Secondary Antibody used were: goat anti-mouse IgG Alexa Fluor 488 A11029 Invitrogen (1:200); goat anti-rabbit IgG Alexa Fluor 488 A11034 Invitrogen (1:200); goat anti-mouse IgG Alexa Fluor 555 A21422 Invitrogen (1:200); goat anti-rat IgG Alexa Fluor 555 A21434 Invitrogen (A21434); goat anti-rat IgG Alexa Fluor 647 150159 Abcam (1:200); and goat anti-mouse IgG Alexa Fluor™ 647 A21236 Invitrogen (1:200).

### *Drosophila* Glioblastoma Model

Glioblastoma (GB) was induced in flies using the established model by Read et al., based on the co-expression of constitutively active forms of dEGFR and dp110 in glial cells. These alterations mimic common mutations observed in GB patients and reproduce key tumor features, including glial overproliferation, invasive behavior, and dysregulation of signalling pathways. Transgene expression was controlled using the temperature-sensitive Gal80ts system: UAS constructs are maintained inactive at 17 °C and are induced upon shifting to 29 °C.

### Mouse models of invasive nervous system cancers

For the neuroblastoma, a transgenic mouse model with Cre-conditional induction of MYCN in dopamine β-hydroxylase-expressing cells (LSL-MYCN;Dbh-iCre) was used to generate MYCN-amplified neuroblastoma (Althoff et al., 2015). Adrenal glands of age-matched healthy control animals were used as controls. At approximately 13 weeks of age, tumors and healthy adrenal glands were freshly dissected, flash-frozen, and stored at -80°C until RNA extraction. For medulloblastoma, a genetically engineered mouse model characterized by constitutive activation of the Hedgehog signalling pathway was used to generate SHH medulloblastoma (SHH MB) tumors. Tumor induction was achieved through tamoxifen-inducible expression of a constitutively active form of Smoothened (SmoM2) in cerebellar granule neuron precursors under the control of the Math1 promoter (Math1-CreER; SmoM2; Math1-GFP), as previously described (Schüller et al. 2008; Tiberi et al. 2014). To induce Cre-mediated recombination, tamoxifen (TAM; Sigma-Aldrich) was freshly prepared and administered at 1 mg/kg via a subcutaneous injection to postnatal day 3 pups. Control animals lacking the oncogenic allele (Math1-CreER; Math1-GFP) were also generated. Five weeks following tamoxifen induction, tumor tissue from four Math1-CreER; SmoM2; Math1-GFP animals were freshly dissected, flash-frozen, and stored at -80°C until RNA extraction. Cerebella from four control animals were collected and processed identically. All experiments were approved by the Animal Welfare and Ethics Body and Direção-Geral da Alimentação e Veterinária (DGAV,Portuguese Authority).

### Medulloblastoma immunohistochemistry

Five weeks after tamoxifen induction, six tumor-bearing animals and two age-matched controls were deeply anaesthetized and transcardially perfused with phosphate-buffered saline. Intact heads were harvested, immersion-fixed in 10% neutral-buffered formalin, decalcified, paraffin-embedded, and sectioned at 4 µm. Sagittal sections through the cranial midline, including the cerebellar vermis, were processed for HPSE1/heparanase immunohistochemistry using anti-Heparanase 1 antibody (Abcam, ab288438). Sections were developed with DAB chromogen and counterstained with hematoxylin. Slides were reviewed by a board-certified veterinary pathologist (TC) using a Leica DM2000 microscope coupled to a Leica MC170 HD camera.

### RNA Isolation and Quantitative RT-PCR

RNA was extracted from fly (20 adult flies) using Direct-zol RNA Microprep (Zymo Research #R2061) according to manufacturer instructions, followed by generation of cDNA with SuperScript III First-Strand Synthesis SuperMix (ThermoFisher). Quantitative PCR was performed with GoTaq qPCR Master Mix on the CFX96 Real-Time System. Expression values were normalized to transcript levels of housekeeping genes.

Mouse tissue was flash-frozen in liquid nitrogen immediately after collection and stored at –80 °C until processing. Total RNA was extracted using the RNeasy® Micro Kit (QIAGEN, Hilden, Germany), according to the manufacturer’s instructions. RNA concentration and purity (A260/280 ratio) were assessed using a NanoDrop 2000 spectrophotometer (Thermo Fisher Scientific, Waltham, MA, USA). Total RNA (100 ng of each sample) was reverse-transcribed into cDNA using the High-Capacity RNA-to-cDNA Kit (Applied Biosystems, Foster City, CA, USA), following the manufacturer’s protocol. Quantitative real-time PCR was performed in technical triplicate on a CFX C1000 Touch Thermal Cycler (Bio-Rad, Hercules, CA, USA) using NZY Supreme qPCR Green Master Mix (NZYTech, Lisbon, Portugal). Gene expression levels were normalized to the geometric mean of three reference genes (GAPDH, B2M, and TBP) and analyzed using the comparative Ct (ΔΔCt) method.

List of primers

*e2fbeta*-Fwd: 5′- GTCATCGAGGACGACCAAGGT-3′

*e2fbeta*-Rv: 5′-TCTTGTTGAAGGCAGCAATG -3′

*hpsl*-Fwd: 5′-TCGTGGGAGATTTTTACTTCGC-3′

*hpsl*-Rv: 5′-CGATTGAACCCCTTCCAAATGA-3′

*Gapdh*-Fwd: 5′-GCCAAAAGGGTCATCATCTC-3′

*Gapdh*-Rv: 5′-CACACCCATCACAAACATGG-3′

*Hpse1*-Fwd: 5′-GGCAATGAGCCCAACAGTTT-3′

*Hpse1*-Rv: 5′-TTGGTAGCGATGCGTCCATT-3’

*Hpse2*-Fwd: 5′-GGGCAAAAGGACGGATTTC-3′

*Hpse2*-Rv: 5′-TGTCCAAGGCAACATCACTC-3′

*B2MF*-Fwd: 5’- TCACCCCCACTGAGACTGAT

*B2MF*-Rv: 5’- TCCCAGTAGACGGTCTTGGG

*TBP*-Fwd: 5’- TCACCCCCACTGAGACTGAT

*TBP*-Rv: 5’- AAATCAACGCAGTTGTGCGTG

Normality Shapiro-Wilk test followed by Welch’s t-test were used for statistical analysis.

### Survival rates and Lifespan curves

Female flies were collected right after eclosion in group of 10 animals per vial. After 3 days of temperature adaptation, a single stab lesion was performed in one optic lobe. Flies were transferred to fresh food vials every 2–3 days, and the number of living and dead flies was recorded at each transfer. Age-matched intact flies were used as controls in all experiments. For survival rate analyses, the percentage of surviving flies per vial was quantified at a fixed time point. In experiments conducted at 25°C, survival was assessed at 40 days, whereas in experiments conducted at 29°C, survival was assessed at 25 days.

For lifespan analyses, Kaplan–Meier survival curves were generated and analyzed using GraphPad Prism. Survival rate and lifespan data were obtained from independent experiments.

### Climbing Assay

The climbing assay was performed essentially as described in Manjila and Hasan 2018. Briefly, 10 flies were transferred to empty vials and allowed to acclimatize for 3 minutes before being tapped to the bottom of the vial. Climbing behavior was recorded, and the climbing index was calculated as the proportion of flies crossing a 10 cm mark within 10 seconds. Three technical replicates were performed, with at least 1 minute of recovery between trials. A baseline climbing assay (Day 0) was performed, and flies were assessed at 1 day and 1 week post injury. For tumor experiments, flies were assessed at baseline and 10 days after tumor induction. The fold change was calculated by normalizing the climbing index of experimental flies to the mean climbing index of the controls at the same timepoint and 2-way Anova test in Graphpad was applied.

### Brain dissociation and FACS

Stab lesions and dissection were performed as previously described. All work surfaces and forceps were cleaned with 70% EtOH and then sprayed with RNase cleaner (Nzytech #MB16001). For OL dissociation, tissue was processed as described in Nagoshi et al., 2010 adapted to OLs. OLs were incubated with 1 µl/OL of activated Papain (Worthington Biochemical Corporation #LK003176) for 25 min at 25 C. Sample was subsequently dissociated using flame-rounded filter tips and passed through a sterile cell strainer (Fisherbrand #22363548). Between 100-400 GFP positive cells from OLs of perma-twin flies were sorted into PCR tubes of 0.5 mL (Axigen® #321-05-051) with 2.5μl Buffer RLT Plus (Qiagen #1053393) by a calibrated BD FACS Aria Fusion (BD Biosciences, US) using a stringent single cell precision mask with a nozzle of 100μm. After recovery, tubes were on ice and immediately frozen at -80°C. For non-injured conditions >120 adult brains of PT flies were dissected to obtain control samples, which were processed within 2h.

### Bulk RNAseq

Total RNA was extracted and processed with the SMART-SEQ2 protocol. RNA integrity was confirmed on an Agilent 2100 Bioanalyzer. Only RNA samples with A260/A280 > 2.0, RIN > 7.0 and undetectable gDNA levels were used for RNA-seq. Libraries were generated with Nextera and sequenced with NextSeq500 Illumina (single-end (SE), 75-base-pair (bp) for 20 million reads). Clean reads were mapped to the Drosophila melanogaster reference genome (release 6 plus ISO1 MT) using STAR 2.7.0a. After sequencing, libraries containing approximately 20 million reads, and no less than 10 million, were generated. Only samples with uniquely mapped reads >71.48% were used for further analyses. Quality control was made in FastQC and then trimmed with Trimmomatic software to remove low scores and sequences <50bp.

### Bioinformatic analyses

Bulk RNA-seq count matrices were analyzed in RStudio® (version 3.6.3). Following quality control, 2-4 samples were retained for downstream analyses. Genes with <10 total counts across all samples were excluded. Count data were analyzed using DESeq2 (version 1.44.0) and normalized using variance stabilizing transformation (VST). Sample quality was assessed by inspecting count distributions, followed by principal component analysis (PCA) and t-distributed stochastic neighbor embedding (t-SNE). Unsupervised clustering using the HDBSCAN algorithm was performed to identify potential outlier samples prior to differential expression analysis.

Differential expression analysis was performed using 1. DESeq2 and 2. limma, performed independently on normalized expression values. P-values were adjusted using the Benjamini–Hochberg method to control the false discovery rate (FDR). Genes were considered differentially expressed (DEGs) if they showed a p-adjusted < 0.05 and an absolute log2 fold change (log2FC) > 1 in at least one of the two methods. For genes identified by both approaches, a consensus DEG list was generated for downstream analyses. Data visualization was performed using ggplot2 and pheatmap. Heatmaps were generated from VST-transformed expression values after row-wise scaling and included DEGs and selected gene sets associated with proliferation, neuroblast proliferation, neuroblast differentiation, and gliogenesis, based on reported and predicted functions in FlyBase. Functional enrichment analyses were performed using clusterProfiler, DOSE, and org.Dm.eg.db. Gene Ontology, KEGG, and Gene Set Enrichment Analysis were performed using DEGs or pre-ranked gene lists with an FDR threshold of 0.05. Functional annotation was additionally performed using DAVID Bioinformatics Resources (2021 update) to complement the biological interpretation of enriched pathways.

### Image analysis: cell number, intensity and volume

In the case of pH3, dad-GFP and *hpsl>RED-stinger,* the number of positive cells per optic lobe (40 μm section) was quantified in Fiji (ImageJ) by manually counting labeled cells that colocalized with DAPI-stained nuclei. pH3 counts were normalized to the non-injured control genotype. For membrane-associated *dpp*>CD8-GFP and HA::DPP signals, quantification was performed similarly, with positive signal identified in the perinuclear region rather than by nuclear colocalization. Dlp staining was quantified by measuring the mean gray value across the entire optic lobe after subtraction of background fluorescence.

The volumes of the Dally::GFP reporter signal and Repo-positive cells were quantified following image segmentation using Labkit (Arzt et al., 2022). Segmented regions were subsequently visualized and analyzed in 3D using the 3D Manager plugin from ImageJ.

For larval glioblastoma, samples were imaged using a Leica TCS SP5 confocal microscope. Quantification of glial cell number was carried out using IMARIS software (Spots and Surface tools).

### Statistics

Statistical analyses were performed with GraphPad Prism (version 8.4.3) software and Rstudio (version 2024.12.0). Statistical tests were chosen based on data distribution and homoscedasticity, with parametric or nonparametric approaches applied as appropriate. Data is shown as mean ± SD. For comparison of injured and non-injured conditions across genotypes, nonparametric Kruskal-Wallis followed by Dunn’s multiple comparison or 2-way ANOVA was performed. For survival assays the statistical test applied were the Long-rank (Mantel-Cox) test and Gehan-Breslow-Wilcoxon test.

Results were considered significant at * p <0.05; ** p <0.01, *** p <0.001, **** p <0.0001.

---

**Figure S1.**
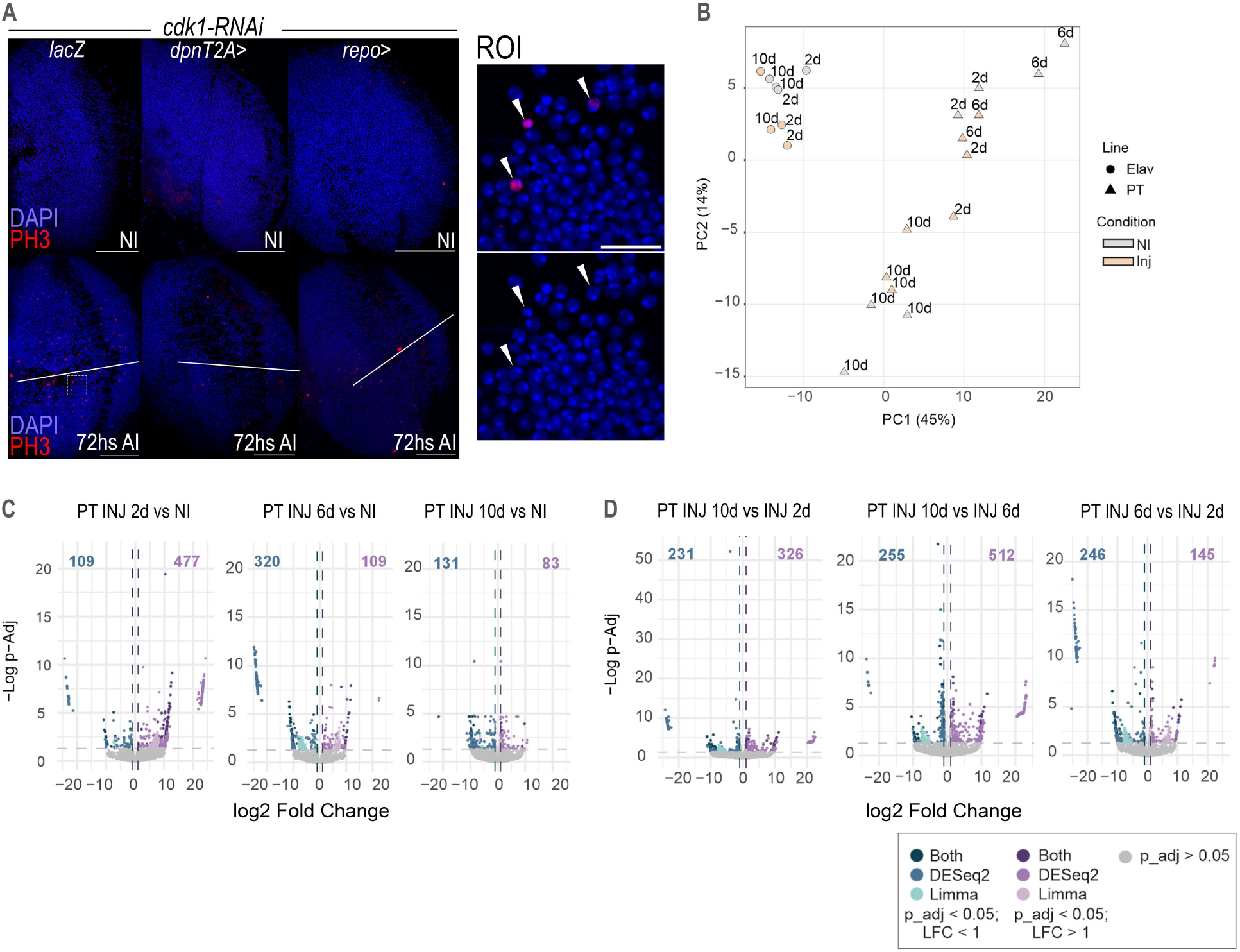
(A) Representative images of pH3 immunostaining in adul optic lobes (OLs), with control (*lacZ*) and *cdk1*-RNAi driven in glia (*repo>*) and neural progentiors (*dpnT2A>*) cells under non-injured (NI) and injured (AI) conditions. Right panels show higher-magnification views of the indicated regions of interest (ROIs). Arrowheads indicate mitotic cells. (B) Principal component analysis (PCA) of RNA-seq datasets obtained from proliferative-tracing (PT) and Elav+ populations under no injury (NI) and injury (INJ) conditions at the indicated time points. (C) Volcano plots showing differentially expressed genes in PT populations isolated from injured (INJ) optic lobes compared with no injury controls (NI) at 2, 6, and 10 days after injury (AI). (D) Volcano plots showing pairwise comparisons between injury time-points within the PT population, illustrating dynamic transcriptional changes during the regenerative response. Scale bars: 50 µm (A), 10 µm (A-ROI)

**Figure S2.**
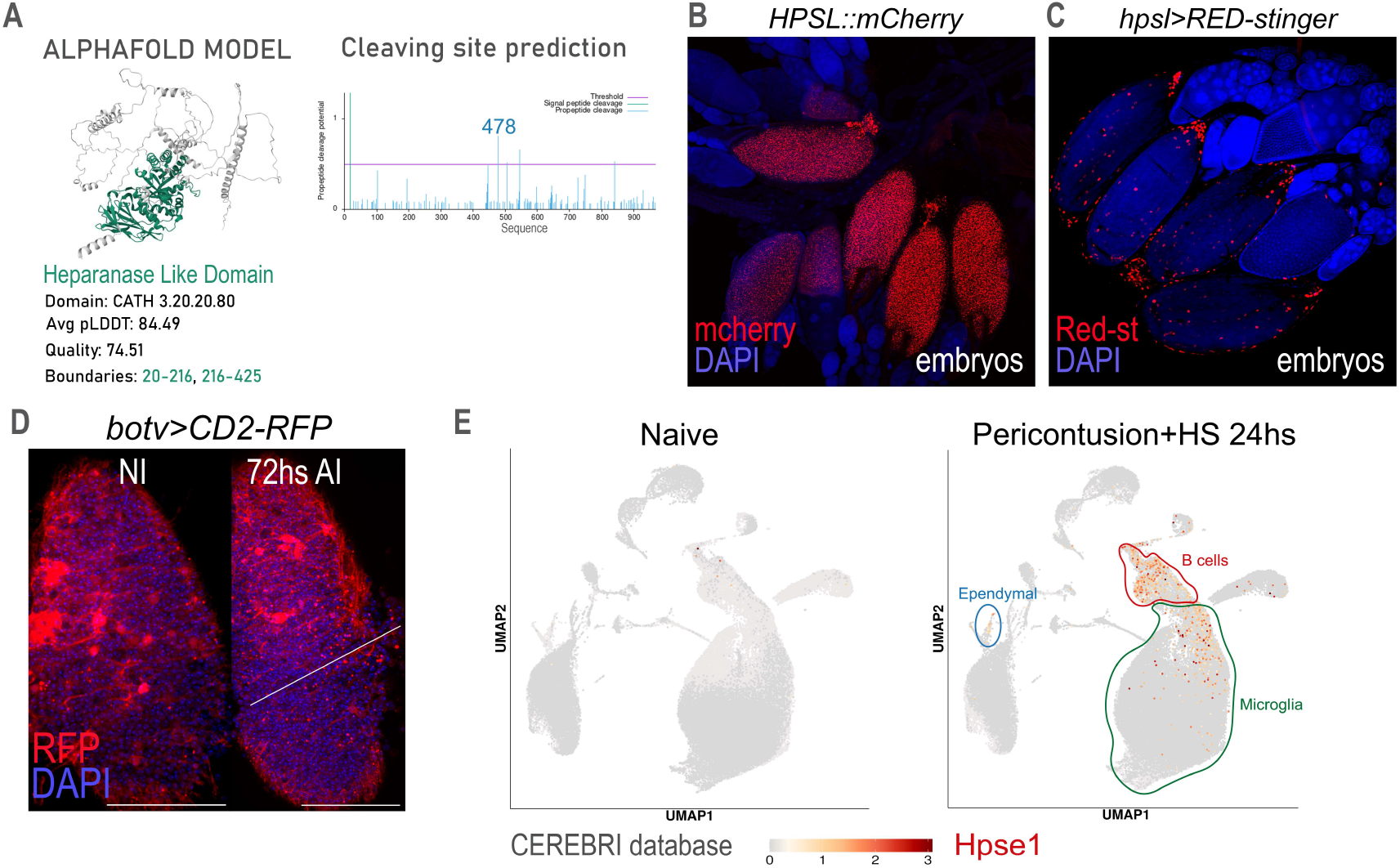
(A) AlphaFold prediction of the HPSL protein structure and identification of a conserved heparanase-like domain. Predicted cleavage sites are indicated. (B) Representative image of endogenous HPSL::mCherry (protein) expression in *Drosophila* embryos. (C) Representative image of the *hpsl* transcriptional reporter (*hpsl>RED-stinger*) in *Drosophila* embryos. (D) Expression pattern of the enzyme Botv (*botv>CD2-RFP*) in control and injury conditions. (E) Visualization of *Hpse1* expression using the CEREBRI single-cell transcriptomic database. UMAP projections from control (naïve) and damage (pericontusion and hemorraghic stroke (HS)) mice brain samples 24 h after injury are shown. Cell populations enriched for Hpse1 expression are indicated. Scale bars: 50 µm (D)

**Figure S3.**
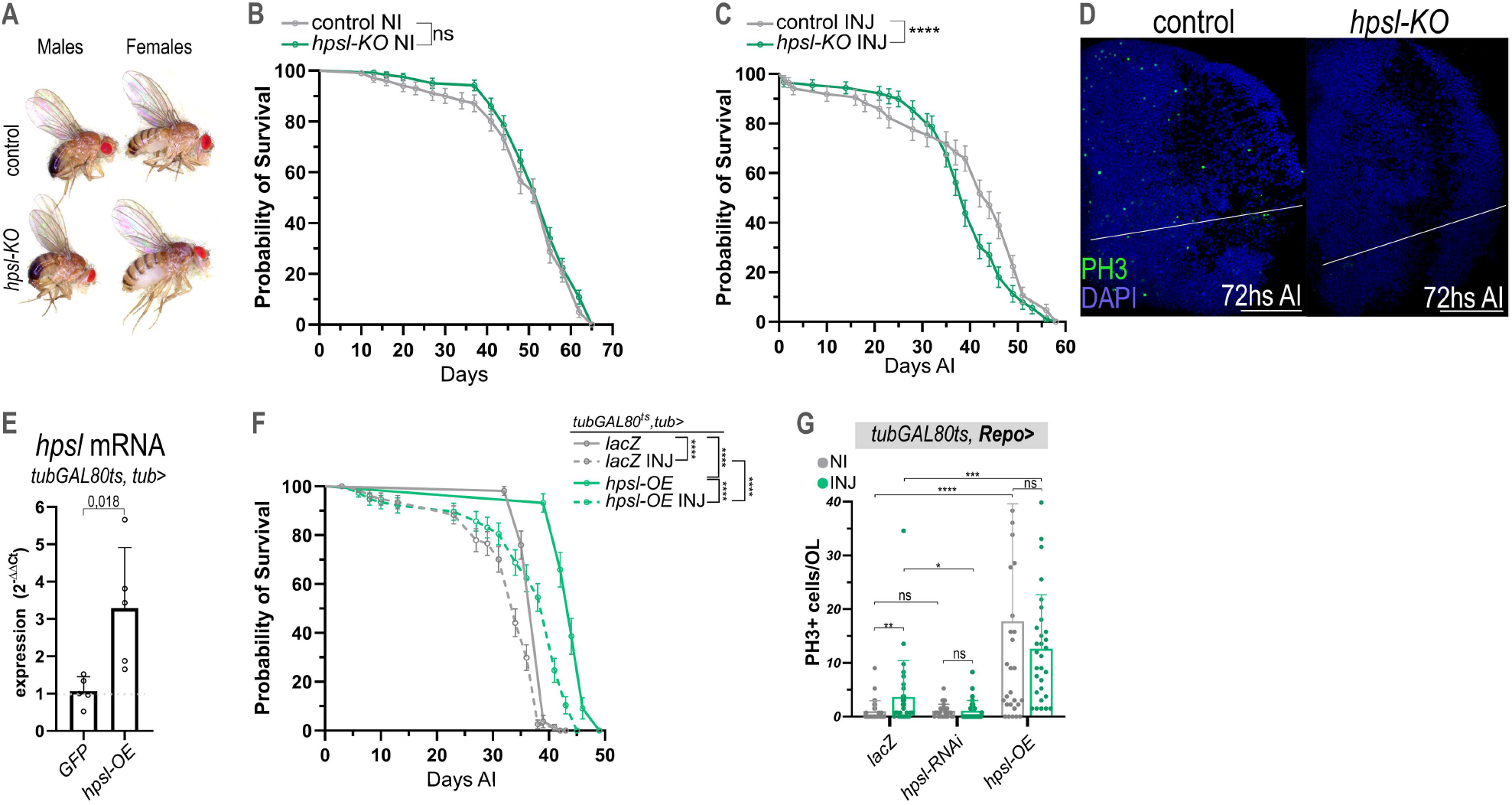
(A) Representative image of adult control and *hpsl-KO* mutant male and female flies. (B) Survival curves of of control and *hpsl*-KO flies without injury (NI). ns (non-significant). (C) Survival curves of control and *hpsl-*KO animals following brain injury (INJ). **** p <0.0001 as determined by Mantel-Cox test. (D) Representative confocal images of optic lobes (OLs) from control and *hpsl-*KO animals at 72 h after injury (AI), stained for the mitotic marker pH3 (green) and DAPI (blue). (E) Relative *hpsl* mRNA expression in adult heads from control (*gfp-OE*) and *hpsl*-OE animals. Means and standard deviation are shown. T-test analysis and p-value as mentioned. (F) Survival curves of non-injured (NI) and injured (INJ) control (*lacZ*) and *hpsl-OE* flies. **** p <0.0001 as determined by Mantel-Cox test. (G) Quantification of pH3+ cells per OL in control (*lacZ*), *hpsl-*RNAi, and *hpsl-*OE animals under NI and INJ conditions. Data is shown as mean ± SD. (G) Quantification of pH3+ cells per OL in control (*lacZ*), *hpsl-*RNAi, and *hpsl-*OE animals under NI and INJ conditions. Data is shown as mean ± SD. Kruskal–Wallis test with Dunn’s multiple comparison correction and p-value as follow: *p <0.05, **p <0.01, ***p <0.001, ****p <0.0001, non-significant (ns). Scale bars: 50 µm (D)

**Figure S4.**
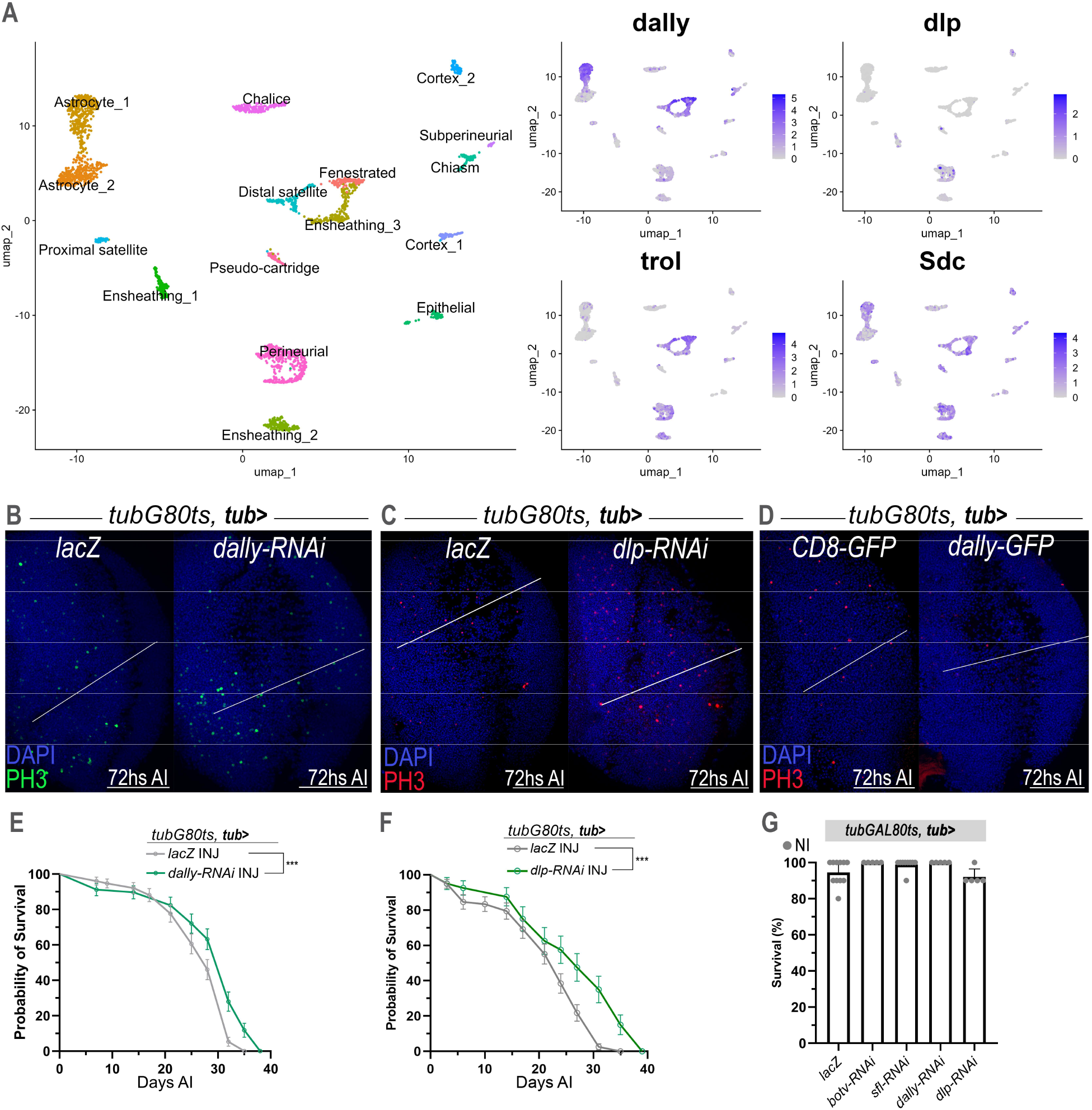
(A) Expression of Heparan Sulfate Proteoglycans (HSPGs) in single-cell transcriptomic dataset from adult optic lobe (OL) as reported by Oezel et. al., 2018 and further analyzed in Lago-Baldaia et al., 2023. (B) Representative images of pH3+ mitotic cells in control (*lacZ*) and *dally-RNAi* OLs 72 h after injury (AI). (C) Images of pH3 immunostaining in control (*lacZ*) and *dlp*-RNAi OLs after injury (AI). (D) Representative images of pH3 immunostaining in control (CD8-GFP) and dally-GFP OLs 72 h AI. (E) Survival curves of injured control (*lacZ*) and *dally*-RNAi animals. *** p <0.001 as assessed by Mantel-Cox. (F) Survival curves of injured control (*lacZ*) and *dlp*-RNAi animals. *** p <0.001 as assessed by Mantel-Cox. (G) Survival of non-injured animals expressing the indicated RNAi constructs. One-way ANOVA with Dunnett’s multiple comparisons. Scale bars: 50µm (B), (C) and (D)

**Figure S5.**
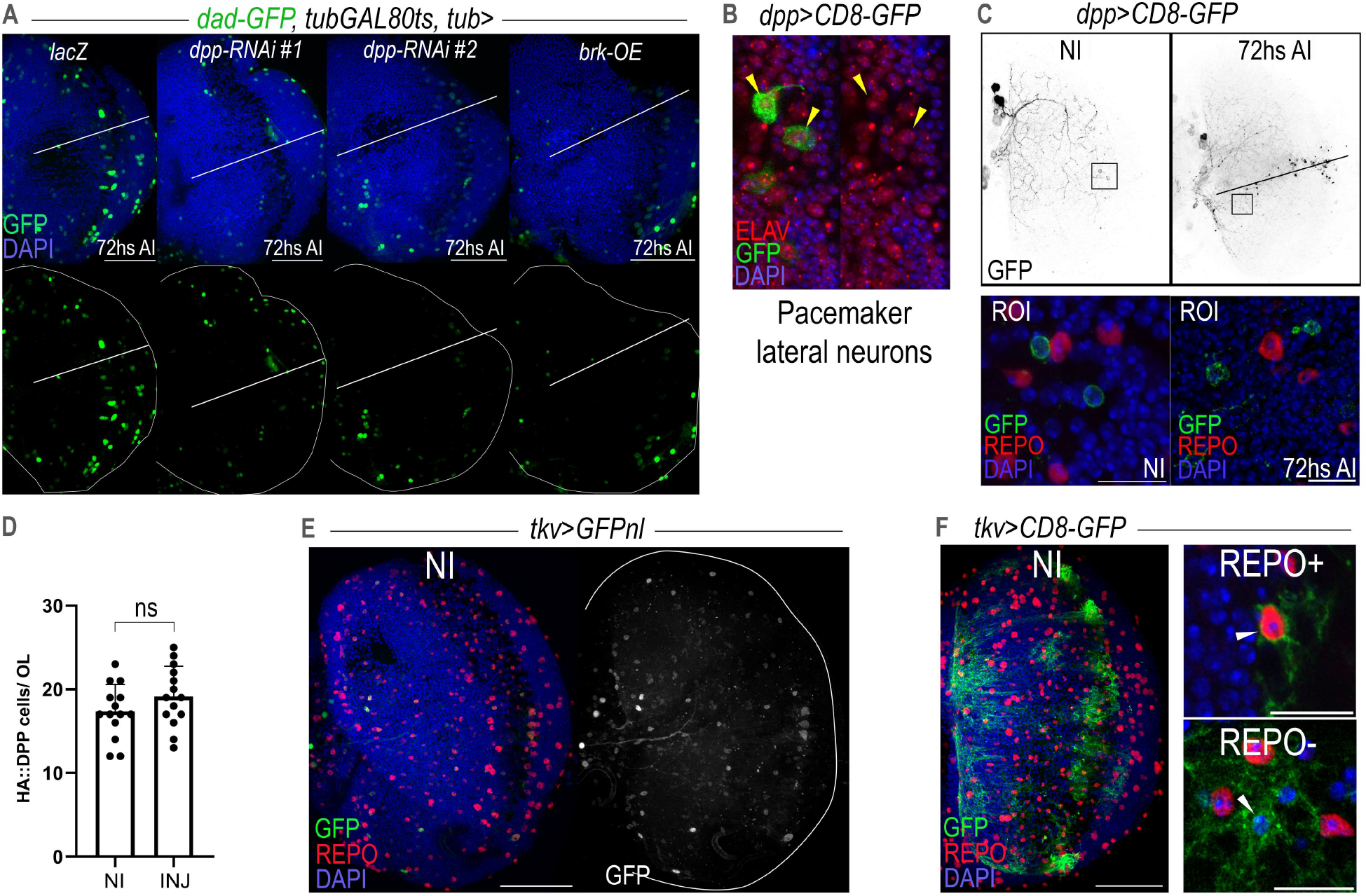
(A) Representative images of dad-GFP expression following knock-down of *dpp* (*dpp-*RNAi) or overexpression of Dpp signalling pathway inhibitor Brk (*brk*-OE). Nuclei are counterstained with DAPI (blue). Lower panels show GFP signal alone. (B) Higher-magnification image of *dpp*>CD8-GFP-positive cells in the region located between the central brain and the optic lobe. Arrowheads indicate representative GFP+ pacemaker lateral neurons (ELAV+, red). (C) Representative images of *dpp*>CD8-GFP expression in non-injured and injured optic lobes (OLs). Lower panels show magnified views of the indicated regions of interest (ROIs). (D) Quantification of HA::DPP-positive cells per OL in no injury (NI) and 72hs after injury (INJ). Data is shown as mean ± SD. Non-significant (n.s.) as assessed by t-test. (E) Representative image of *tkv>GFPnl* expression in the adult OL. (F) Representative images of *tkv>CD8-GFP* expression in the adult OL. To the right, colocalization with both, Repo+ glial cells and Repo- cells are indicated with arrowheads. Scale bars: 10µm (B) and ROIs in (C) and (F); and 50µm (A), (E) and (F)

**Figure S6.**
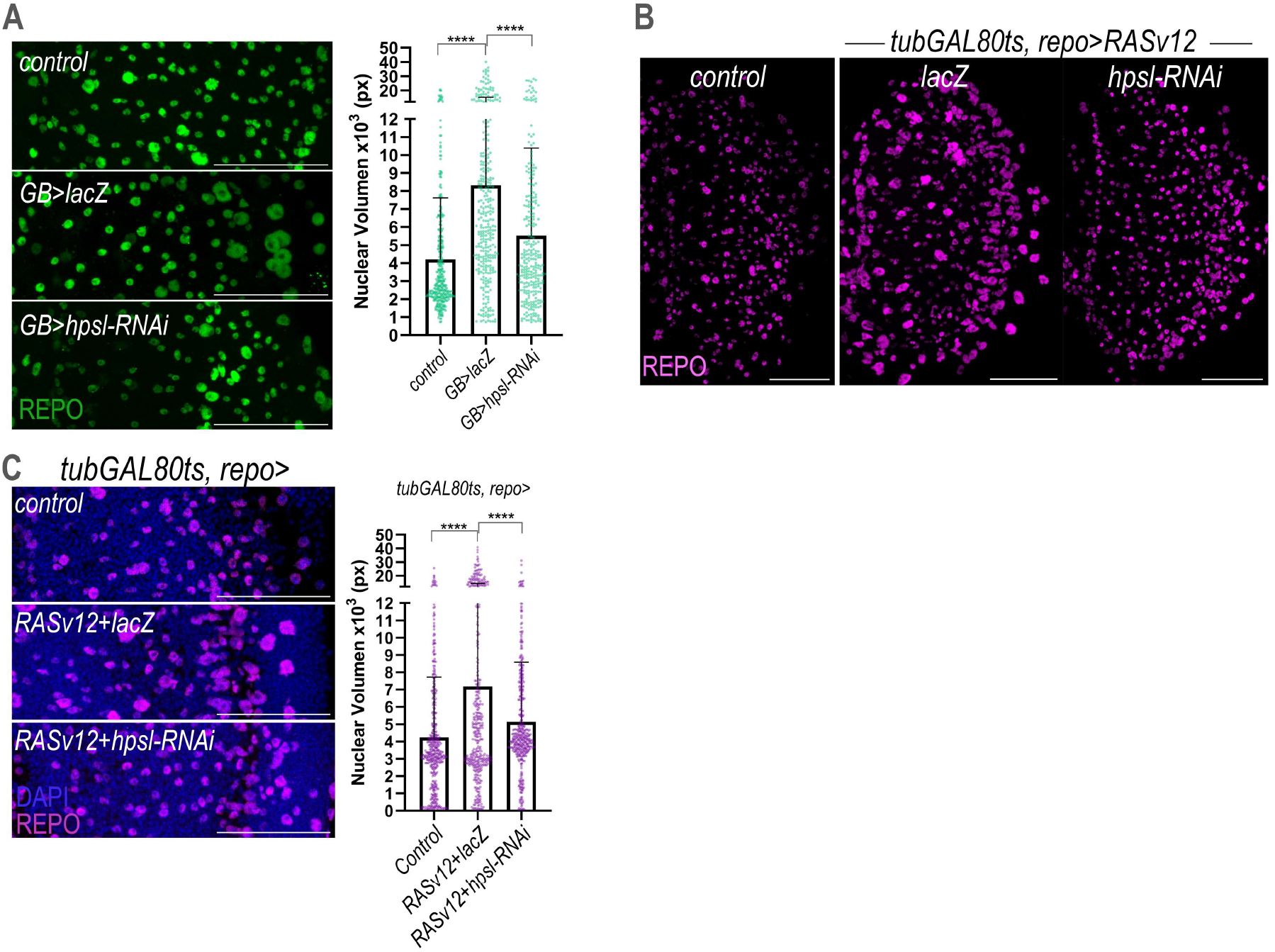
(A) Images of glial-specific expression of control (*repo>lacZ*) and Glioblastoma (GB>) cells activated in adult briains, combined with *lacZ* or *hpsl*-RNAi constructs. REPO staining is shown in green. Quantification of REPO+ glial nuclear volume is shown on the right. Data is shown as mean ± SD. ****p <0.0001 as assessed by Mann–Whitney–Wilcoxon. (B) Representative images of adult optic lobes (OLs) from control animals and of a glial RasV12-driven model (*tubGAL80ts, repo>RasV12*), combined with (*lacZ*) or *hpsl*-RNAi constructs. REPO staining is shown in magenta. (C) Representative images of adult OLs from control flies and flies with adult onset RASV12-expression in glia (*tubGAL80ts, repo*>), in the presence of control *(lacZ*) or *hpsl*-RNAi constructs. DAPI and REPO staining (magenta) are shown. Quantification of REPO+ glial nuclear volume is shown on the right. Data is shown as mean ± SD. ****p <0.0001 as assessed by Mann–Whitney–Wilcoxon. Scale bars: 50 µm (A), (B) and (C).

## Notes

### Competing Interest Statement

The authors have declared no competing interest.

